# Biochemically diverse CRISPR-Cas9 orthologs

**DOI:** 10.1101/2020.04.29.066654

**Authors:** Giedrius Gasiunas, Joshua K. Young, Tautvydas Karvelis, Darius Kazlauskas, Tomas Urbaitis, Monika Jasnauskaite, Mantvyda Grusyte, Sushmitha Paulraj, Po-Hao Wang, Zhenglin Hou, Shane K. Dooley, Mark Cigan, Clara Alarcon, N. Doane Chilcoat, Greta Bigelyte, Jennifer L. Curcuru, Megumu Mabuchi, Zhiyi Sun, Ryan T. Fuchs, Ezra Schildkraut, Peter R. Weigele, William E. Jack, G. Brett Robb, Česlovas Venclovas, Virginijus Siksnys

## Abstract

CRISPR-Cas9 nucleases are abundant in microbes. To explore this largely uncharacterized diversity, we applied cell-free biochemical screens to rapidly assess the protospacer adjacent motif (PAM) and guide RNA (gRNA) requirements of novel Cas9 proteins. This approach permitted the characterization of 79 Cas9 orthologs with at least 7 distinct classes of gRNAs and 50 different PAM sequence requirements. PAM recognition spanned the entire spectrum of T-, A-, C-, and G-rich nucleotides ranging from simple di-nucleotide recognition to complex sequence strings longer than 4. Computational analyses indicated that most of this diversity came from 4 groups of interrelated sequences providing new insight into Cas9 evolution and efforts to engineer PAM recognition. A subset of Cas9 orthologs were purified and their activities examined further exposing additional biochemical diversity. This constituted both narrow and broad ranges of temperature dependence, staggered-end DNA target cleavage, and a requirement for longer stretches of homology between gRNA and DNA target to function robustly. In all, the diverse collection of Cas9 orthologs presented here sheds light on Cas9 evolution and provides a rich source of PAM recognition and other potentially desirable properties that may be mined to expand the genome editing toolbox with new RNA-programmable nucleases.

## Introduction

The Cas9 protein from type II CRISPR (clustered regularly interspaced short palindromic repeats)-Cas (CRISPR-associated) antiviral defence systems have been repurposed as a robust genome editing tool (reviewed in^1–5^). Target DNA specificity is accomplished with small non-coding RNAs that through direct base pairing guides Cas9 to its DNA target^6,7^. In addition to guide RNA (gRNA) recognition, a sequence motif, termed the PAM (protospacer adjacent motif), is required for the initiation of Cas9-guide RNA target binding and cleavage^6,7^. Easily reprogrammed to recognize new DNA sequences, it has been widely adopted for use in a multitude of applications to edit genomic DNA, modulate gene expression, visualize genetic loci, or detect targets *in vitro*^8–15^. To date, just a handful of variants are used for these applications^16–18^ with the *Streptococcus pyogenes* (Spy) Cas9 being used most widely^2^.

Since Cas9 can be programmed to target DNA sites by altering the spacer sequence of the gRNA, recognition of the PAM becomes a constraint which restricts the sequence space targetable by Cas9. This is further limited by the requirement for careful site selection to minimize off-target binding and cleavage based on the tolerance for mismatches in the gRNA-PAM-target complex^19–23^. This constraint becomes particularly evident in therapeutic applications where even rare genome alterations resulting from off-targets are undesirable or when targeting more structurally complex plant genomes^24,25^. Moreover, these restraints impact the use of Cas9 for homology-directed repair (HDR), template-free editing, base editing, or prime editing applications, where the outcome is reliant on the proximity of the desired change to the target sequence^26–32^. Furthermore, the biochemical and physical characteristics of Cas9, producing predominantly blunt-end DNA target cleavage^6,7^, slow substrate release^33^, low frequency of recurrent target site cleavage^34^, gRNA exchangeability^35^, temperature dependence^36^, and size^37^ may also be unfavourable for its varied applications.

While Spy Cas9 targeting constraints are beginning to be addressed through structure guided rational design^38–40^ and directed evolution approaches^41–44^, we show that the diversity provided by naturally occurring orthologs may offer unique insight and opportunities for improvement of this powerful tool. First, we determined the gRNA and PAM requirements for 79 phylogenetically distinct Cas9s of various sizes without the need for protein purification or extensive computational analyses^45^. In doing so, we identified extraordinary diversity in Cas9 PAM and gRNA requirements. This extended the number of unique classes of gRNAs from four to seven and revealed T-, A-, C-, and G-rich PAM recognition that varied in length from one to more than four nucleotides. Interestingly, analysis of the PAM interacting (PI) domain indicated that much of this variation was derived from just 4 related groups. Finally, additional biochemical studies revealed new diversity that may further extend application. This included differences in temperature and spacer length requirements as well as variation in the pattern of doublestranded DNA target cleavage.

## Results

### Cas9 ortholog selection

To systematically sample diversity, 47 orthologs were chosen from most of the 10 major clades of a Cas9 evolutionary tree (Figure 1A and Supplementary Table S1). Clades giving rise to previously characterized proteins that were active in eukaryotic cells were mined at a rate of approximately 20% while all others were surveyed at a rate of approximately 10%. To enrich for proteins with robust biochemical activity and thermostability, an additional 32 orthologs were selected based on their physiochemical properties (e.g. predicted secondary structure and isoelectric point), classification as a type II-A subtype^35,46^, and affiliation with a thermophilic host organism (Supplementary Table S2). Sequence length variation of our collection matched that found in naturally occurring orthologs and ranged from ~1,000 to ~1,600 residues with a bimodal distribution focused around sizes of ~1100 and ~1375 amino acids (Supplementary Figure S1). Furthermore, sequence alignments of those selected showed extraordinary variation, altogether, differing by as much as 93% (Supplementary Table S1).

**Figure 1.**
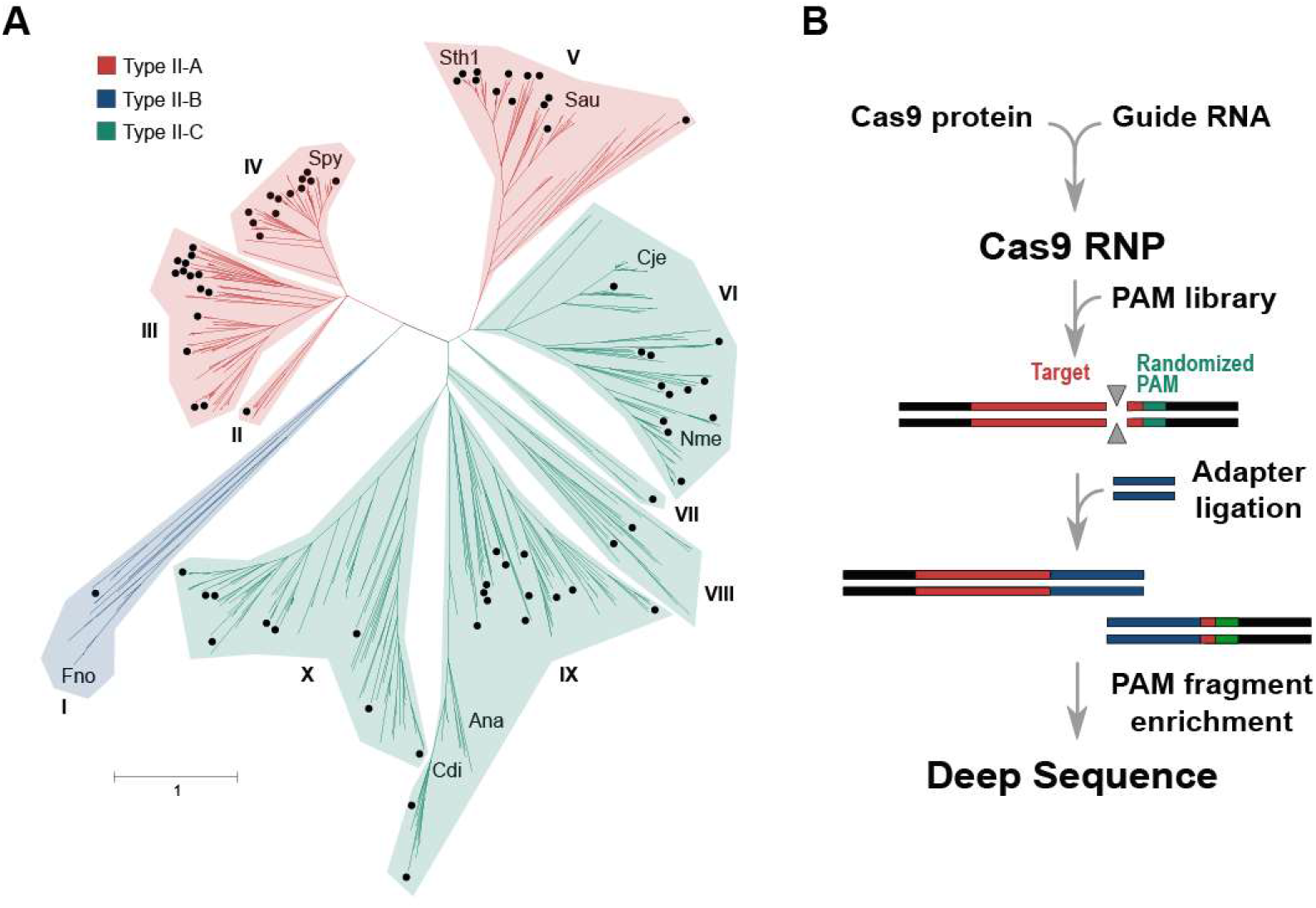
Cas9 diversity and characterization approach. **(A)** Phylogenetic representation of the diversity provided by Cas9 orthologs. Type II-A, B, and C systems are color-coded, red, blue, and green, respectively. Distinct phylogenetic clades are numbered I-X. Those selected for study are indicated with a black dot. Cas9s whose structure has been determined are also designated. **(B)** Biochemical approach used to directly capture target cleavage and assess PAM recognition. Experiments were assembled using Cas9 protein produced by IVT.

### Guide RNA requirements

In all instances, Cas9 gRNAs, the crRNA (CRISPR RNA) and tracrRNA (trans-activating CRISPR RNA), were identified near the *cas9* gene, however, spatial positioning as well as the transcriptional orientation varied greatly among the systems characterized (Supplementary Figure S2). In general, these features were conserved among orthologs belonging to a particular phylogenetic clade (Supplementary Figure S2). Most CRISPR repeats were ~36 bp length, however, longer repeats (45-50 bp), associated with orthologs from clade X, were also identified (Supplementary Table S2). Computational analyses comparing co-variant models (CMs) based on sequence and secondary structure homology among the characterized tracrRNAs showed 7 distinct clusters (Figure 2). For some Cas9 orthologs, the tracrRNA self-clustered or demonstrated weak similarity to other CMs (Figure 2). In these cases, it was not assigned to a particular group. In general, clusters were tightly associated with a particular Cas9 phylogenetic clade, although, exceptions were noted (Figure 2). Examination of the sgRNA modules (repeat:anti-repeat duplex, nexus, and 3 ‘ hairpin-like folds^47,48^ were also typically conserved among related Cas9 proteins (Supplementary Figure S3). For example, the sgRNA solutions for almost all members of clade IV resembled that belonging to Spy Cas9 and comprised a bulge in the repeat:anti-repeat duplex, a short nexus-like stem-loop, and two hairpins followed by a poly-U sequence at the 3’end^47,48^. Analogous structures were observed in the sgRNAs of Cas9 proteins belonging to clades VIII and X. However, in clade X sgRNAs, the repeat:anti-repeat duplexes were typically fully complementary and didn’t form repeat:anti-repeat bulges. Members of clade V contained the shortest sgRNAs, and reminiscent of the sgRNA from *Streptococcus aureus* (Sau) Cas9, these contained only 2 hairpins (nexus-like followed by a larger fold) following the repeat:anti-repeat duplex (Supplementary Figure S3). In contrast, sgRNAs associated with clades III, VI, and IX orthologs displayed longer, more complex and diverse structures. These included a variety of differences in stem length, presence of bulges, and different spacing between sgRNA modules (Supplementary Figure S3). In addition, it was more difficult to reliably identify a Rho-independent-like terminator at the end of some tracrRNA encoding regions for these clades.

**Figure 2.**
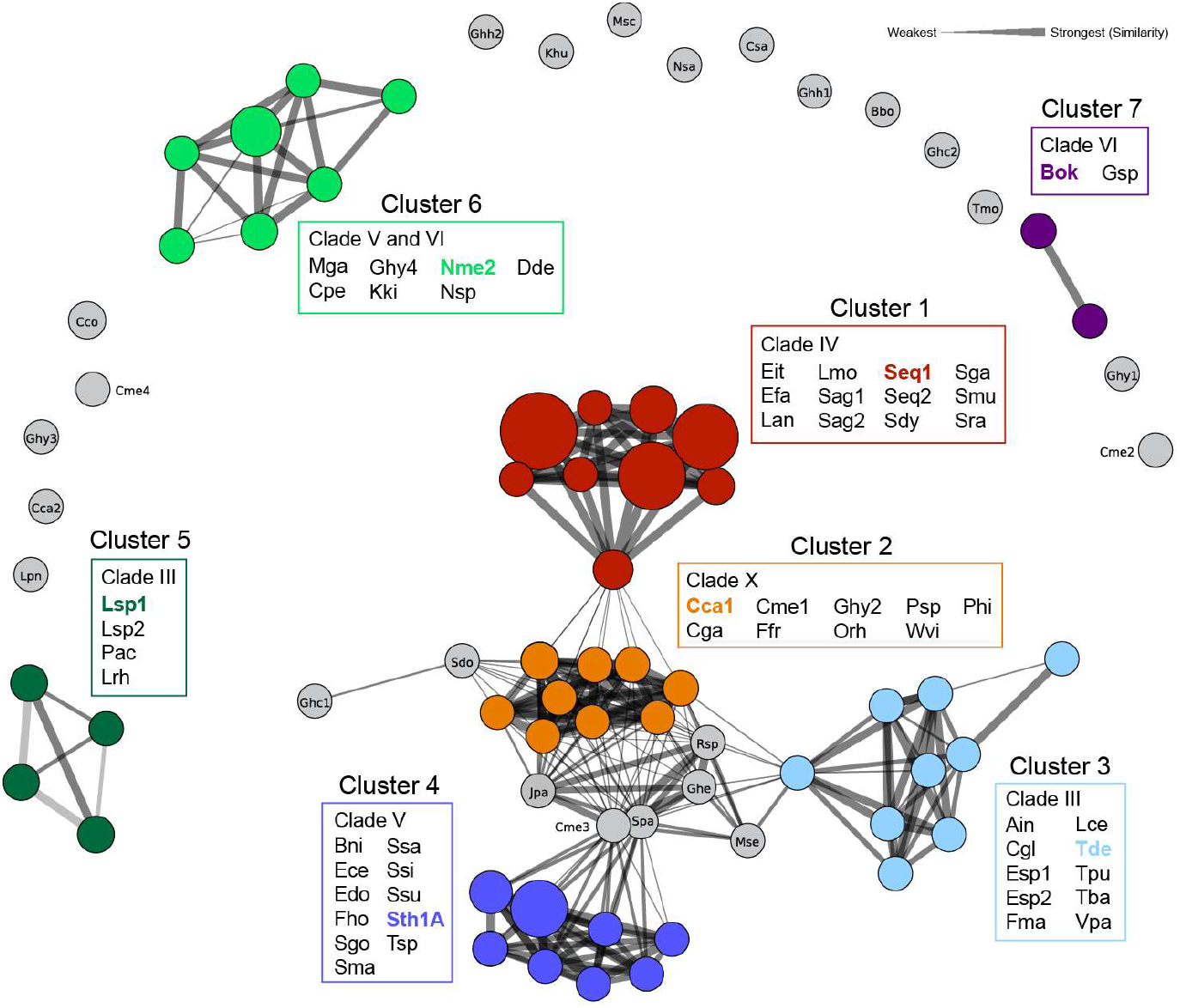
Cas9 tracrRNA sequence and secondary structure similarity. Circles are scaled based on the number of sequences belonging to each covariance model (CM) and colored according to the designated cluster. The width of the connecting lines indicates the percentage of similarity or relatedness among CMs. Representative tracrRNAs from each cluster are indicated with the associated color. CMs not assigned to a cluster are in grey.

### PAM recognition by orthologous Cas9s

To rapidly survey the target recognition properties of new Cas9s, we employed a cell-free *in vitro* translation (IVT) method similar to that described previously (Figure 1B)^45,49^. Since PAM recognition is dependent on the concentration of Cas9-guide RNA complex^50^, crude IVT RNP mixtures were diluted (10^1^ to 10^3^ in 10-fold increments) and tested for their ability to support cleavage when combined with a plasmid library containing a randomized PAM region adjacent to a Cas9 target site. The greatest dilution supporting cleavage activity was then used as a baseline for PAM recognition. To confirm the accuracy of our approach, Cas9 PAM recognition was examined using purified components as described previously^50^. This was done for Spy, S. *thermophilus* CRISPR3 (Sth3), and S. *thermophilus* CRISPR3 (Sth1) Cas9s, whose PAM was determined previously^50^ and for 11 orthologs from our collection. As shown in Supplementary Figure S4A & B, there was a nearly perfect agreement between the approaches. Additionally, to examine the propensity for PAM recognition to extend beyond position 7 (the length of randomization in our PAM library), the spacer targeting the PAM library was also shifted 5’ by 1, 2 or 3 nts for Cas9 orthologs that exhibited PAM preferences at positions 6 or 7 and lacked PAM requirements in the first, second or third positions. This permitted PAM identification to be extended to 8, 9, or 10 bp, respectively. Of the 20 Cas9s that were tested (Supplementary Table S2), only 5, all belonging to phylogenetic clade VI (Figure 1A & 2), had PAM recognition that continued beyond the 7^th^ position. Surprisingly, PAM preferences at the 8^th^ position were always an A residue similar to the previously characterized *Brevibacillus laterosporus* (Blat)^50^ and *Geobacillus stearothermophilus* (Geo)^51^ Cas9 proteins. Altogether, we found that long PAMs extending beyond the 7^th^ position were not widespread and only abundant in one family of orthologs belonging to clade VI.

The Cas9 orthologs characterized (Supplementary Table S2) with our IVT-based approach demonstrated significant divergence in PAM recognition. Indeed, we identified nucleases with novel PAM requirements that varied in composition both in sequence and length. Among these were proteins with PAM recognition that could be generally sub-divided into A-, T-, and C-rich PAM recognition in addition to the G-rich PAM typical of the Spy Cas9 protein (Figure 3). PAMs composed of multiple residues of a single base pair while present (e.g. Efa, Nme2, Rsp, Ssi, and Ssu) were rare but notably enriched in Clade IV (e.g Efa) to which Spy Cas9 belongs, and in Clade VII (Ssi, Ssu) (Figure 1A, Figure 3, and Supplementary Table S3). In general, Cas9s with composite PAM recognition containing at least two different base pairs were more abundant (Figure 3). The length of PAM recognition also varied between 1-4 base pairs or more with most orthologs exhibiting recognition at three or more positions. Additionally, many proteins exhibited seemingly degenerate PAM recognition. Typically, this resulted in a strong requirement for at least one base pair in combination with positions that accepted more than one (typically two) base pairs (e.g. Lan, Mse, Nsa, and Sma).

**Figure 3.**
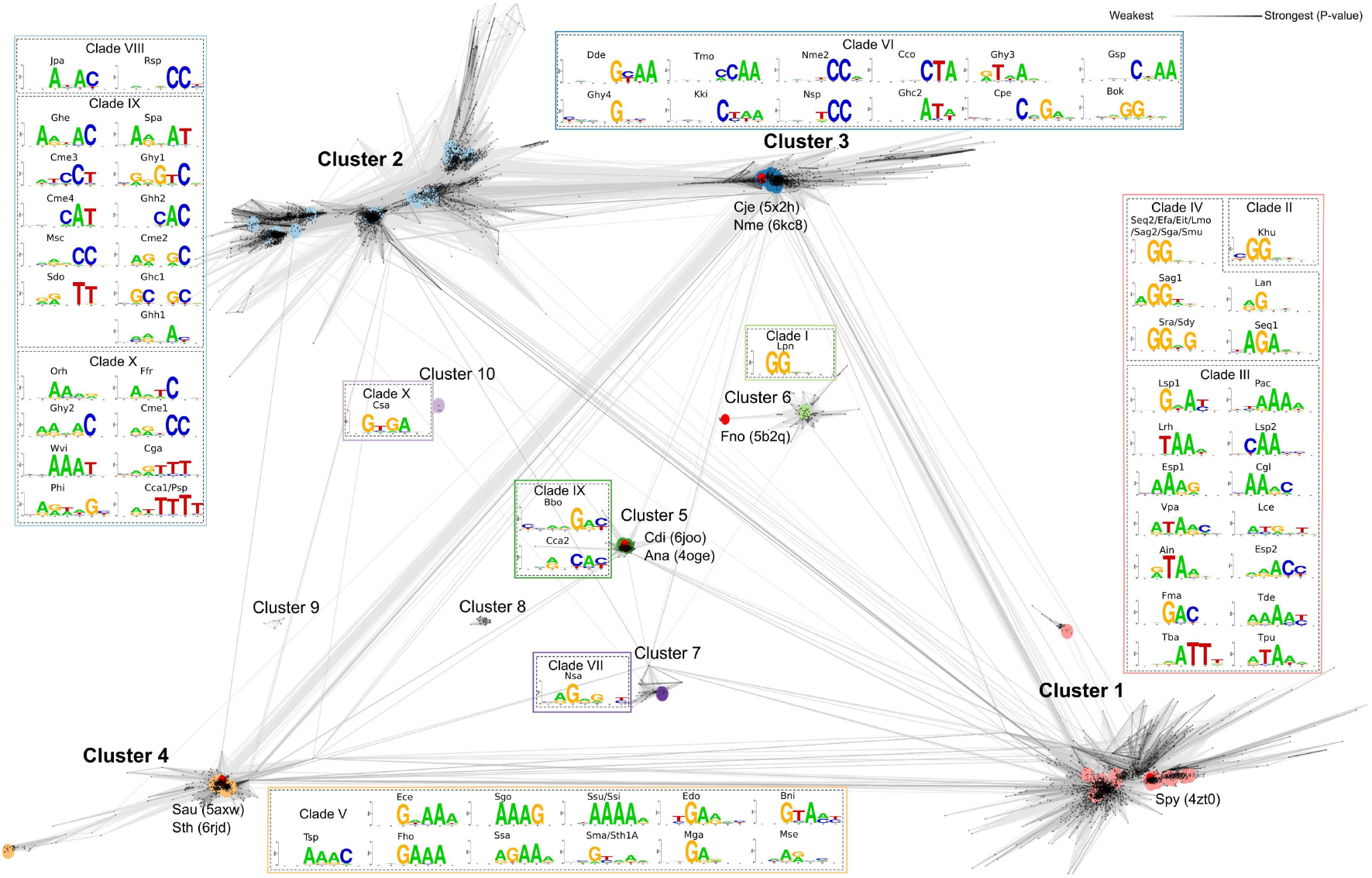
Cas9 PAM Interacting (PI) domain similarity. Cas9 PI domains clustered by their pairwise sequence similarity. Lines connect sequences with P-value ≤ 1e-11. Line shading corresponds to P-values according to the scale in the top-right corner (light and long lines connect distantly related sequences). Major clusters are shown in bold. Cluster 1 was so named to emphasize that it contains first experimentally characterized Cas9. Clusters 2 to 10 were named beginning from the one with the most members. Sequences having known structures are marked red, their PDB code is shown in parentheses.

### Diversity and taxonomic distribution of Cas9 PAM interacting domains

The extreme diversity of experimentally determined PAM sequence requirements prompted us to evaluate the sequence relationship of Cas9 PAM interacting (PI) domains. To do so, we extracted the PI regions from the characterized orthologs and used them as queries for iterative searches against non-redundant collections of microbial proteins (see methods). In all, 9,161 sequences having non-identical PI domains were found (Supplementary Table S4). Sequences were next clustered based on their pairwise similarity leading to identification of ten clusters (Figure 3). Clusters 1-4 were the largest and contained 93% of all sequences recovered while clusters 7-10 were considerably smaller and were comprised of 4 to 37 sequences (Figure 3, Supplementary Table S4). Sequence searches with HHpred^52^ showed that most clusters were distantly related to each other (Figure 3 and Supplementary Figure S5, A-C) with an exception being cluster 10 that did not reveal significant similarity to any other group (Supplementary Figure S5, D). In general, PI domain similarity could be correlated with the major phylogenetic branches of the Cas9 tree (Figure 1A and Figure 3). For example, Cas9s belonging to clades II, III, and IV grouped into cluster 1 (Figure 3). Additionally, phylogenetic analysis of clusters 1-6 also suggested that similar PI domains usually resulted in similar PAM recognition (Supplementary Figure S6 A, B, D), however, sequence diversity and length varied greatly even among members of the same group (Supplementary Figure S6C). Closer examination of the Cas9s belonging to cluster 1 further highlighted that even within similar PI architectures sequence composition varied considerably with conservation being the lowest in the PI domain relative to the rest of the Cas9 protein (Supplementary Figure S7).

Although clusters shared amino acid sequence similarity (Figure 3), their taxonomic distribution differed. While PI domains from cluster 2 were mainly found in *Bacteroidetes* and *Alphaproteobacteria* (Supplementary Figure S6B), cluster 3 was more likely to come from *Betaproteobacteria, Epsilonproteobacteria*, and *Firmicutes* (*Bacilli* and *Clostridia*) (Supplementary Figure S6C). Sequences from clusters 1 and 4 were usually found in *Firmicutes* (Supplementary Figure S6 A & D) while clusters 5 and 6 were specific to *Actinobacteria* and *Proteobacteria*, respectively (Supplementary Figure S5 E).

### Evaluation of Cas9 ortholog biochemical activity

Fifty-two Cas9 orthologs from our collection were selected for additional characterization using purified components. Primary selection criteria included simple PAM recognition (≤3 bp) (where possible) while maintaining diversity in phylogenetic distribution and protein size. It was previously reported that Sau and Geo Cas9 proteins require a longer spacer to function robustly^16,51^. Therefore, we designed sgRNAs with two different spacer lengths, 20 and 24 nt, for each ortholog and examined their influence on Cas9 cleavage activity *in vitro*. Exceptions to this included Efa, Lpn and Cme4 where a single spacer length of either 20 or 22 nt was tested. As shown in Supplementary Figure S8, most orthologs worked best with a 20 nt spacer similar to Spy Cas9 when evaluated across a panel of 5 different buffers, however, six orthologs, Cga, Cca1, Orh, Tmo, Nsa, and Ghh1 Cas9, required a spacer length of greater than 20 nt to effectively cut their DNA target. In all, 46 out of 52 produced double-stranded DNA target cleavage activity greater than 25% under the conditions examined.

The thermal stability of thirty-eight orthologs showing robust target cleavage were next predicted using nano differential scanning fluorimetry (nanoDSF). 36 of 38 proteins showed a melting temperature of >37°C confirming stability under standard *in vitro* enzymatic reaction conditions. Interestingly, five orthologs had melting temperatures >50°C suggesting thermostability (Supplementary Figure S9). These included Cme2, Cme4, Ghy1, Esp1 and Nsa Cas9.

To corroborate nanoDSF predictions, DNA target cleavage was next measured in reactions at temperatures ranging from 10°C to 70°C. In all, Cas9 orthologs displayed a wide spectrum of temperature dependencies including both narrow and broad ranges of activity (Figure 4A). Consistent with thermal unfolding analysis, Cme2, Esp1, Nsa, Ain, Cme3, and Sth1A, were active at temperatures greater than 50°C with Nsa, isolated from the deep-sea hydrothermal vent chimney bacterium, *Nitratifractor salsuginis*^53^, remaining active at temperatures greater than 60°C (Figure 4B). Additionally, one ortholog, Ssa, retained 95% of its cleavage activity at 10°C (Figure 4A). We also observed that 5 Cas9 orthologs (Cme2, Cme4, Nsp, Khu, and Fma) retained less than 25% activity at reaction temperatures of 25°C or below.

**Figure 4.**
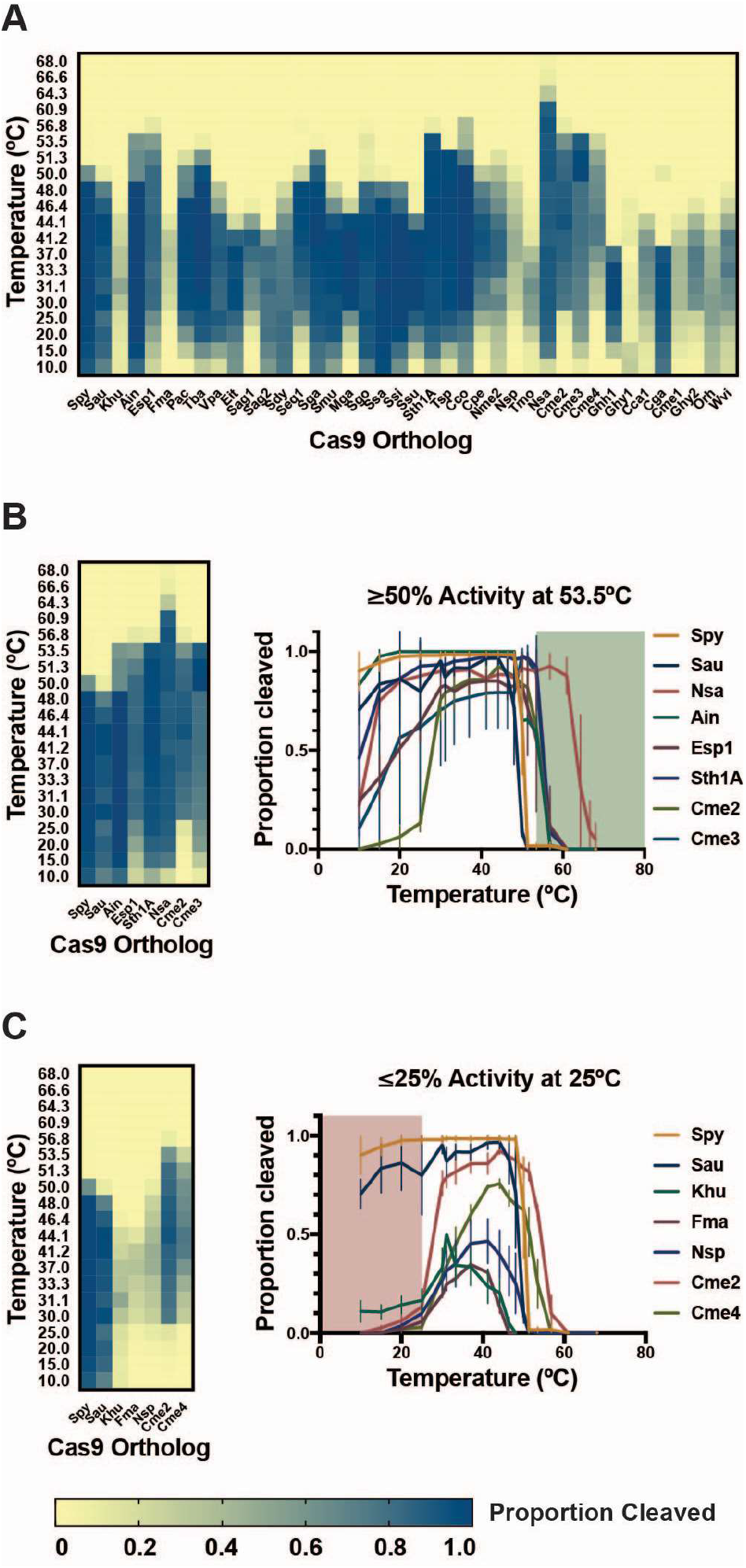
Activity of Cas9 orthologs at varying temperatures. The cleavage activity of Cas9 orthologs was measured using *in vitro* DNA cleavage assays using fluorophore-labeled dsDNA substrates. Cleaved fragments were quantitated and are represented in a heat map (A) showing overall activity at temperatures ranging from 10°C to 68°C. (B) Cas9 orthologs with activity at elevated temperatures. *In vitro* DNA cleavage activity for a subset of Cas9 orthologs with >50% activity at 53°C is summarized in a heat map and plotted as proportion of DNA substrate cleaved at varied temperature. Points represent the mean +/- SEM of at least 3 independent experiments. (C) Cas9 orthologs with reduced activity at room temperature. *In vitro* DNA cleavage activity for a subset of Cas9 orthologs with <25% activity at 25°C is summarized in a heat map and plotted as a proportion of DNA substrate cleaved at varied temperature. Points represent the mean +/- SEM of at least 3 independent experiments.

### Target DNA cleavage by Cas9 orthologs

To characterize the termini resulting from Cas9 DNA cleavage, we developed a method which allows both termini resulting from target cleavage to be captured simultaneously in a deep sequencing read (Supplementary Figure S10). To validate the approach, we examined the cleavage positions for restriction endonucleases HhaI, FspI, and HinP1I, and for Spy Cas9. As shown in Figure 5A, these enzymes generated defined cut-sites, either blunt-ended or staggered as expected. This can be contrasted with Spy Cas9 target cleavage that, depending on target site, generated either blunt-end cuts, 1 nt 5’ staggered termini due to RuvC post-cleavage trimming as observed previously^54^, or a mixture of both (Supplementary Figure S11). In all, as averaged across 5 targets, Spy Cas9 generated predominantly blunt-ended DNA target cleavage as reported previously^7^ (Figure 5B). Analysis of Sau Cas9 target cleavage produced almost entirely blunt-ended products as shown earlier ^54^ (Figure 5B and S11 and Supplementary Table S5).

**Figure 5.**
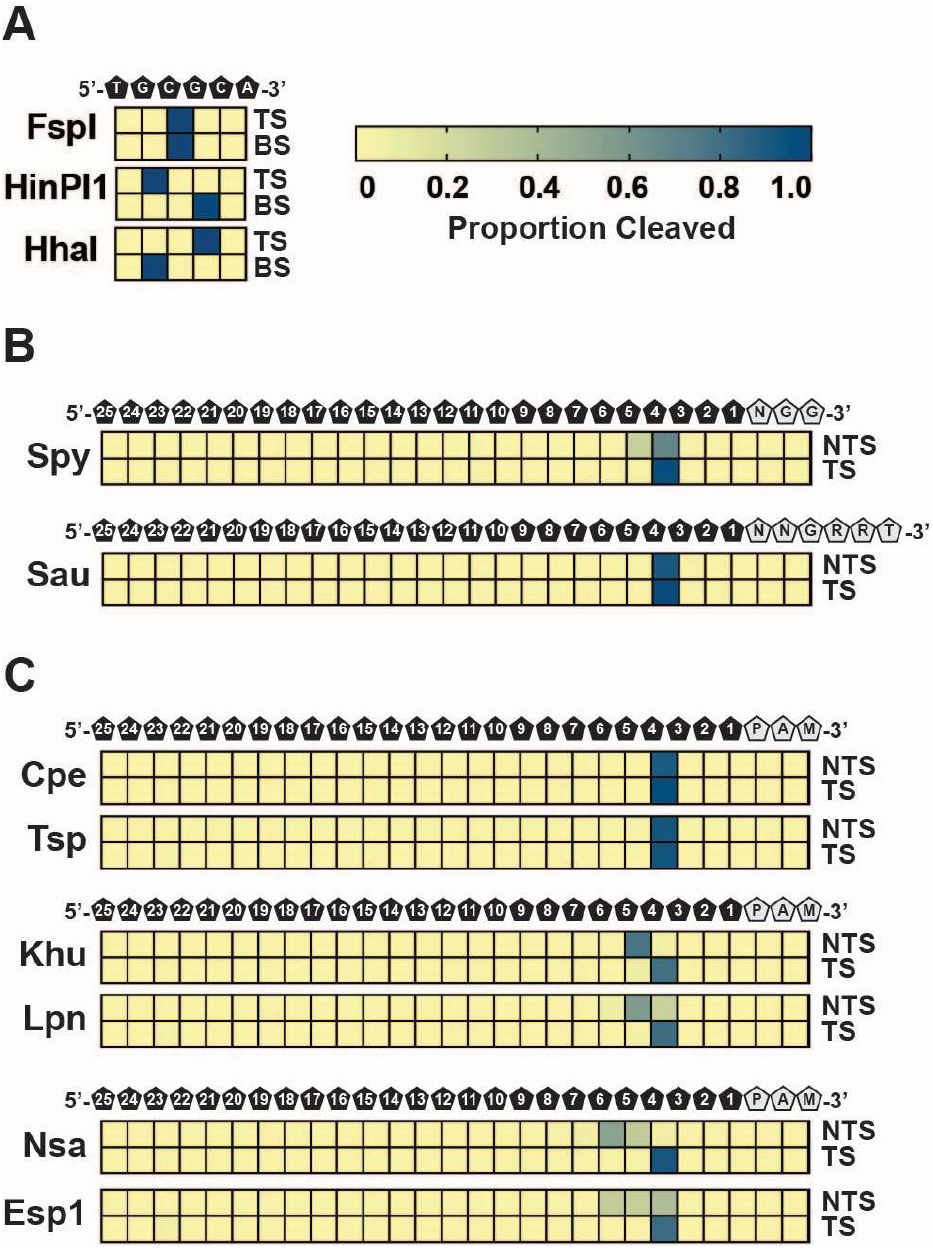
Target DNA cleavage patterns produced by Cas9 orthologs. Cleavage sites and resultant dsDNA ends are depicted as heatmaps that show the proportion of cleaved ends recovered by DNA sequencing at each position of a target DNA. Intensity of the blue color indicates the proportion of mapped cleavage ends. (A) Control digests using restriction enzymes recover 5’-overhangs, 3’-overhangs, and blunt ends. TS indicates top strand; BS indicates bottom strand. (B) SpyCas9 and SauCas9 cleaved DNA ends. Heatmaps represent mapped cleavage ends as the averages at each position in 5 different dsDNA targets. Position of the DNA bases and PAM sequences is depicted above the heatmaps. NTS indicates non-target strand; TS indicates target strand. (C) Blunt and staggered-end cleavage. Examples of blunt, one base 5’-overhang staggered cleavage, and multiple base 5’-overhang cleavage are depicted as heatmaps that show the proportion of cleaved ends as the averages at each position in 5 different dsDNA targets. Position of the DNA bases and PAM sequences is depicted above the heatmaps. NTS indicates non-target strand; TS indicates target strand.

We next evaluated the target cleavage pattern for 19 orthologs from our collection. Some Cas9s produced blunt-ended termini like Sau Cas9 (e.g. Cpe and Tsp (Fig 4C)) while others, depending on the target site, produced either blunt-ended or 1 nt 5’-overhang termini similarly to Spy Cas9 (e.g. Sag1 and Seq1 (Supplementary Table S5)). Some orthologs, as averaged across 5 different target sites, consistently generated 5’-overhang termini varying between 1 or more nts (e.g. Khu, Lpn, Nsa, and Esp1) (Figure 5C and Supplementary Table S5). In these cases, the non-target strand tended to terminate at multiple positions suggesting variation in the positioning of or post-cleavage trimming by the RuvC domain while the target strand was cleaved predominantly between the 3^rd^ and 4^th^ positions of the protospacer (Figure 5C and Supplementary Table S5).

## Discussion

We identified Cas9s with novel G-rich, C-rich, A-rich, and T-rich PAM recognition of varying compositions, altogether, greatly expanding the sequence space targetable by Cas9. The observed diversity in PAM length was also striking with the majority of orthologs recognizing PAMs greater than 2 bp. This difference may be important for genome editing applications as orthologs with longer PAM recognition (≥3 bp) may afford higher specificity^16,17,55^. Additionally, phylogenetic and clustering analyses revealed that the PI domain was not always congruent with the rest of the protein. For example, conservation of the PI domain among related Spy Cas9 proteins was 1.4 times lower relative to the N-terminal portion (Supplementary Figure S7). In some cases, these differences were even greater as was noted for orthologs from *Neisseria meningitidis* (Nme) where PI domains only shared 52% identical residues while the rest of the protein was nearly the same (98% identity)^56^ or in clade X where sequences belonged to three different homology clusters (Figure 3; clusters 7, 5 and 10). Moreover, sequence variation from clusters 1, 3 and 4 when compared with the structures of Spy (4zto), Nme (6kc8), Sau (5axw) and Sth (6rjd) could be modelled into just a single PI domain architecture (Supplementary Figure S12). Altogether, these observations could in part be explained by the uncoupling of PI domain evolution from the rest of protein indicating that it is under selective pressure to diversify perhaps in response to PAM-based phage escape strategies as described previously^57–59^. Additionally, they suggest that the Cas9 PI domain is extraordinarily flexible and can be engineered to recognize a wide variety of sequence motifs encompassing the full spectrum of DNA nucleotides.

The gRNAs from our collection were in general conserved between related Cas9 proteins, although, new and diverse gRNA structures were observed. In general, they could be classified into 7 groups based on tracrRNA sequence and structural homology and visual inspection of sgRNA modules as exemplified by Seq1 (Spy-like), Cca1, Tde, Sth1A, Lsp1, Nme2, and Gsp (Figure 2 and Supplementary Figure S3). Altogether, this may warrant the expansion of the number of discrete non-cross reactive Cas9 and sgRNA combinations from 4 to 7 or more pending future studies^35,60^. This finding is important for orthogonal genome editing approaches where simultaneous, yet disparate activities are required at different sites^60,61^.

Finally, an in-depth evaluation of DNA cleavage activity of the Cas9 nucleases described here exposed additional differences among orthologs. These included a wide range of temperature dependencies. Of particular interest was Cme2 Cas9, which was only robustly active from ~30°C to 55°C suggesting the possibility of temperature-controlled DNA search and modification. Additionally, the DNA cleavage activity at different temperatures for Nsa and Ssa Cas9s suggested they could be harnessed for use in thermo-or psychrophiles, respectively. Furthermore, we characterized orthologous Cas9 nucleases with different, and potentially advantageous properties compared to those generally prescribed to Spy Cas9. These included variation in the termini resulting from target cleavage as well as a preference for a longer tract of gRNA and DNA target site homology.

## Methods

### Identification and phylogeny of Cas9 orthologs

Type II Cas9 endonucleases were identified by searching for the presence of an array of CRISPRs using PILER-CR^62^ Following identification, the DNA sequences surrounding the CRISPR array (about 15 kb 5’ and 3’ of the CRISPR array) were examined for the presence of open-reading frames (ORFs) encoding proteins greater than 750 amino acids. Next, to identify *cas* genes encoding Cas9 orthologs, multiple sequence alignment of sequences from a diverse collection of Cas9 proteins was performed using MUSCLE^63^ and then used to build profile hidden Markov models (HMMs) for Cas9 sub-families as described previously^35^ using HMMER^64,65^. The resulting HMMs were then utilized to search protein sequences translated from the *cas* ORFs for the presence of genes with homology to Cas9. Alternatively, Cas9 orthologs and the metagenomic sequence encoding them were obtained from publicly available datasets through the Joint Genome Institute’s Integrated Microbial Genomes & Metagenomes resource (IMG/M): https://img.jgi.doe.gov/cgi-bin/m/main.cgi^66^ Only proteins containing the key HNH and RuvC nucleolytic domains and catalytic residues defining a type II Cas9 protein^67^ were selected (Supplementary Table S6). Through phylogenetic analyses (MEGA7^68^) Cas9 proteins were then parsed into distinct families and representative members of each group used to select orthologs for characterization. To place our collection in context with previously described Cas9 orthologs, a phylogenetic tree was built using type II-A, -B, and -C representatives^69^ and those we selected for characterization using MEGA7^68^ employing Neighbor-Joining^70^ and Poisson correction^71^ methods.

### Engineering single guide RNA solutions

The trans-activating CRISPR RNA (tracrRNA) essential for CRISPR RNA (crRNA) maturation^72^ and Cas9 directed target site cleavage in type II systems^7,73^ was identified by searching for a region in the vicinity of the *cas9* gene, the anti-repeat, which may base-pair with the CRISPR repeat and was distinct from the CRISPR array(s). Once identified, the possible transcriptional directions of the putative tracrRNAs for each new system were established by examining the secondary structures using UNAfold^74^ and VARNA^75^ and possible termination signals present in RNA versions corresponding to the sense and anti-sense transcription scenarios surrounding the anti-repeat. Based on the likely transcriptional direction of the tracrRNA and CRISPR array, single guide RNAs (sgRNAs), representing a fusion of the CRISPR RNA (crRNA) and tracrRNA^7^, were designed. For each ortholog, this was accomplished by linking 16 nt of the crRNA repeat to the complementary sequence of the tracrRNA anti-repeat by a 4 nt GAAA loop similar to that described previously for Spy Cas9^7^. All repeat, tracrRNA sequences and sgRNA solutions are listed in Supplementary Table S2.

### Computational analysis of Cas9 tracrRNAs

BLAST (with parameters to optimize finding short sequences in highly-repetitive regions (-task blastn_short-dust no))^76^ was used to identify sequences homologous to the 79 identified tracrRNAs. The resulting collection of identified sequences were grouped using CD-HIT^77^ at a 90% sequence similarity threshold. The resulting clusters were filtered to remove groups that did not contain at least one of the 79 reference tracrRNA sequences. Next, sequence homology and secondary structure models were constructed for each group using MAFFT^78^ and RNAalifold^79^, respectively. Both models were then used to search for sequence/structural homology in the full set of reference and BLAST-identified sequences using the RNA structure search tools in the Infernal v1.1 software suite^80^. Structural overlap between clusters was then generated by comparing the results of each covariance model (CM). To graph the relationship among tracrRNAs, vertices were first added for each representative CM (sequences with both shared secondary structure predictions and at least 90% sequence similarity). If two vertices shared a CM, they were connected with a line weighted by the percent similarity between shared vertices (percent similarity=(# of shared sequences)/(min(# found by model 1,# found by model 2))).

### Production of single guide RNAs

All single guide RNA (sgRNA) molecules used in this study were synthesized by *in vitro* transcription using HiScribe™ T7 Quick High Yield RNA Synthesis Kits (New England Biolabs), or transcribed directly in the *in vitro* translation (IVT) reaction. Templates for sgRNA transcription were generated by PCR amplifying synthesized fragments (IDT and Genscript) or by annealing a T7 primer oligo to a single stranded template oligonucleotide. Transcribed RNA products were treated with DNaseI (New England Biolabs) to remove DNA templates and cleaned up with Monarch RNA Cleanup Kit (50μg) (New England Biolabs) and eluted in nuclease-free water. RNA concentration and purity were measured by NanoDrop spectrophotometry and RNA integrity was visualized by SYBR™ Gold staining of reaction products separated on Novex TBE-Urea 15% denaturing polyacrylamide gels with 0.5x TBE (Tris borate EDTA) buffer.

### PAM library cleavage using *in vitro* translation

Cas9 was produced by IVT using either a continuous exchange 1-Step Human Coupled IVT Kit (Thermo Fisher Scientific) or a PURExpress bacterial IVT kit (New England Biolabs), following the manufacturer’s recommended protocol similar to that described previously^45^. Plasmid DNA encoding human or *E. coli* codon optimized Cas9s were generated for use as templates for IVT reactions. Synthetic DNA fragments were synthesized by Genscript, Inc. and Twist Bioscience and assembled by NEBuilder HiFi DNA Assembly kit (New England Biolabs) into pT7-N-His-GST (Thermo Fisher Scientific) or pET28a (EMD Millipore). Following IVT, 20 μl of supernatant containing soluble Cas9 protein was mixed with RiboLock RNase Inhibitor (40 U; Thermo Fisher Scientific) and 2 μg of T7 *in vitro* transcribed sgRNA and incubated for 15 min. at room temperature. Alternatively, the sgRNA was transcribed directly in the IVT kit by supplying a DNA template containing a T7 promoter and sequence encoding the respective sgRNA. In this situation, 0.5 μg of plasmid encoding the *cas9* gene and a 100-fold molar excess of sgRNA template was added to the IVT reaction mix. 10 μl (or series of ten-fold dilutions) of the resulting Cas9-sgRNA ribonucleoprotein (RNP) complex were then combined with 1 μg of the 7 bp randomized PAM library described previously^50^ in a 100 μl reaction buffer (10 mM Tris-HCl pH 7.5 at 37°C, 100 mM NaCl, 10 mM MgCl2, 1 mM DTT) and incubated for 60 min. at 37°C. Finally, cleaved library fragments were captured by adapter ligation, enriched for by PCR amplification, and deep sequenced as described previously^50^.

### Identification of PAM preferences

PAM sequences that supported double-stranded DNA target cleavage were captured by adapter ligation, enriched for by PCR amplification, and deep sequenced as described earlier^45,50,81^. Bias in the bp composition at each position within the randomized PAM library was first adjusted to that in the starting library by normalization ((Treatment Frequency)/((Control Frequency)/(Average Control Frequency))). Then, PAM preferences were quantified using position frequency matrices (PFMs)^82^ and displayed as a WebLogo^83^. Analyses were limited to the top 10% most frequent PAMs to reduce the impact of background noise resulting from non-specific cleavage coming from other components in the IVT mixtures.

### Computational analysis of Cas9 PAM interacting domains

The Cas9 orthologs characterized here were aligned using MAFFT^78^. Their PAM interacting (PI) regions corresponding to the C-terminal domain of *Streptococcus pyogenes* Cas9 (4ZT0_A:1090-1365) were extracted and used as queries for two iterations of PSI-BLAST^84^ search against the NCBI NR protein collection, UniRef100 and MGnify^85^ databases. Hits were extracted, filtered to 80% identity using CD-HIT ^77^ and clustered with CLANS^86^. Resulting networks were visually inspected and clusters identified. For groups larger than 150 sequences (Supplementary Table S4), a phylogenetic analysis was performed recovering sequences which were filtered out during the previous step and removing identical ones. Next, multiple sequence alignments were performed for clusters 1 to 6 using MAFFT (options: “--ep 0.123 --maxiterate 20 --localpair”) and regions with gaps removed with trimAL^87^(option: “-gt 0.01”). Lengths of the resulting alignments varied from 359 to 652 residues in clusters 2 and 3, respectively. Phylogenetic trees were generated using IQtree^88^ with auto model selection and 1000 fast bootstrap (options: “-alrt 1000 -bb 1000”).

### Cas9 expression and purification

Spy, S. *thermophilus* CRISPR3 (Sth3), and S. *thermophilus* CRISPR3 (Sth1) Cas9 proteins were expressed and purified as described previously^50^. Other orthologs were first *E. coli* codon optimized and cloned into the pET28 vector, to yield constructs encoding fusion proteins comprising a C-terminal 6-His-tag. In some instances, sequences encoding nuclear localization sequences (SV40 origin) were incorporated onto the 5’ and 3’ ends of the *cas9* gene. The expression analysis was then performed in different *E. coli* strains (NiCo21(DE3), T7 Express lysY/Iq, NEB® Express Iq) under various growth conditions (media, temperature, induction) and detected by SDS-PAGE analysis. Optimised conditions then were chosen for flask scale purification. Cells were disrupted by sonication. The supernatant was loaded onto HiTrap DEAE Sepharose (GE Healthcare), followed by subsequent purification on Ni2^+^-charged HiTrap chelating HP column (GE Healthcare) and HiTrap Heparin HP (GE Healthcare) columns. Purified Cas9 proteins were stored at −20°C in 20 mM Tris-HCl, pH 7.5, 500 mM KCl, 1 mM EDTA, 1 mM DTT, and 50% (v/v) glycerol.

### Evaluation of protospacer cleavage patterns

To capture protospacer cleavage patterns with single molecule resolution, we developed a minicircle double stranded (ds) DNA substrate that allows both ends of target cleavage to be captured in a single Illumina sequence read. First, 124 nt oligonucleotides (IDT) (see Supplementary Table S7) were circularized using with CircLigase™ single stranded (ss) DNA Ligase (Lucigen) according the manufacturers suggestion. Circularized ssDNA was next purified and concentrated using a Monarch® PCR & DNA Cleanup Kit (NEB). 20 pmol of the purified product was then incubated with 25 pmol of a complementary primer in 1X T4 DNA ligase buffer (NEB) supplemented with 40 μM dNTPs. To allow the primer to anneal, the reaction was then heated to 65°C for 30 seconds followed by a decrease in temperature to 25°C at a rate of 0.2°C/second. 6 units of T4 DNA polymerase and 400 units of T4 DNA ligase (NEB) were then added and the reaction was incubated at 12°C for 1 hour to allow second strand synthesis. Following purification with a Monarch® PCR & DNA Cleanup Kit and elution into 1X

CutSmart^®^ buffer (NEB) containing 1 mM ATP, 15 units of Exonuclease V (RecBCD; NEB) and T5 exonuclease (NEB) were added to the sample and incubated at 37°C for 45 min. 0.04 units of Proteinase K (NEB) was then added and the sample was incubated at 25°C for 15 min. prior to purification with a Monarch^®^ PCR & DNA Cleanup Kit. After elution, the yield of circular double-stranded DNA was assessed using an Agilent 2100 Bioanalyzer.

For minicircle digestion, Cas9 RNPs were formed by incubating 1 pmol of sgRNA with 0.5 pmol of Cas9 protein in 1X NEBuffer™ 3.1 or 2.1 (NEB) at room temperature for 10 min. 0.1 pmol of circular dsDNA substrate was added, samples were incubated for 15 min at 37°C and then each 20 μl reaction was quenched by the addition of 5 μl of 0.16 M EDTA. Reactions were concentrated and purified with a Monarch^®^ PCR & DNA Cleanup Kit and the entire 8 μl of eluted product was used as a substrate for Illumina sequencing library construction using a NEBNext^®^ Ultra™ II DNA Library Prep Kit for Illumina^®^ (NEB) and the protocol provided with the kit. 15 cycles of PCR were used to add the Illumina priming sequences and index barcodes and then the concentration of each reaction was assessed on an Agilent 2100 Bioanalyzer. Libraries were pooled and sequenced on either an Illumina NovaSeq or NextSeq instrument with 2 × 150 paired-end sequencing runs. Cleavage sites were then mapped using custom scripts and visualized as heatmaps (representing proportion) cleaved using GraphPad Prism 8.

### *In vitro* cleavage assays for determining optimal buffer, temperature, and spacer length

First, DNA substrates containing a canonical PAM for each ortholog were amplified from HEK293T genomic DNA by PCR using primers corresponding to WTAP and RUNX1. Forward primers were labelled with 5’-FAM and 5’-ROX for WTAP and RUNX1, respectively. Reverse primers were unlabelled. 515 and 605 bp PCR products for WTAP and RUNX1, respectively, were then purified with a Monarch^®^ PCR & DNA Cleanup Kit (5μg) (NEB T1030S) and DNA concentration and purity measured by NanoDrop™ spectrophotometry (ThermoFisher). Purified Cas9 protein was then diluted to 1μM in dilution buffer (300mM NaCl, 20mM Tris, pH7.5) and stored on ice. Next, sgRNAs (Supplementary Table S2 and S7) were diluted to 2 μM in nuclease-free water. Cas9 and sgRNA were then combined in a 2:1 sgRNA:Cas9 molar ratio in reaction buffer at room temperature for 10 min. Substrate was added next at a Cas9:sgRNA:DNA ratio of 10:20:1 and incubated for 30 min. For buffer optimization and spacer length preference experiments, 1X NEBuffers 1.1, 2.1, 3.1, or CutSmart (NEB B7200S) were used as reaction buffers and incubations took place at 37°C. For thermoactivity experiments, reactions were performed in NEBuffer 3.1. Here, RNPs were initially formed at room temperature and then transferred to a thermal cycler pre-heated or cooled to the various assay temperatures prior to DNA substrate addition. 10x DNA substrate (100nM) was separately equilibrated at the designated temperature prior to being added to the RNP containing reaction tubes. Reactions were quenched by adding SDS to 0.8% (v/v) and 80 mU Proteinase K (NEB P8107S). Cleavage products were diluted 4X in nuclease-free water and subjected to capillary electrophoresis (CE) to quantify the extent of cleavage^89^. The fraction of substrate cleaved at each temperature was then visualized as heatmaps, using GraphPad Prism 8.

### Cas9 protein thermal stability

Purified Cas9 proteins were diluted in 300 mM NaCl, 20 mM Tris, pH7.5 to 5-10 μM at room temperature. 10μL of the diluted protein was loaded into NanoDSF Grade Standard Capillaries (NanoTemper) and melting temperatures were determined using a Prometheus NT4.8 NanoDSF instrument according to the manufacturer’s instruction. Temperature was increased from 20°C to 80°C at the rate of 1°C/min. Inflection points of melting curves are reported as the Tm^90^

## Data Availability

Raw deep sequencing data that support PAM and cleavage pattern determination for Cas9 orthologs were deposited at XXXX under Accession numbers XXXX. All other relevant data are available from the corresponding authors on reasonable request.

## Accession Numbers

## Acknowledgement

We thank Migle Stitilyte from CasZyme for preparation of the sgRNA templates.

## Conributions

G.G., J.K.Y.’ M.C., C.A., N.D.C., W.E.J., E.S., G.B.R. and V.S. designed research; G.G., J.K.Y., T.K., D.K., T.U., M.J., M.G., S.P., P.W., Z.H., S.K.D., G.B., J.L.C., M.M., Z.S., R.F.T., and P.R.W. performed research, and G.G., J.K.Y., C.A., N.D.C., W.E.J., E.S., G.B.R., C.V. and V.S. analyzed data. G.G., J.K.Y., G.B.R. and V.S. wrote the paper. All authors read and approved the final manuscript.

## Supplementary Information

Supplementary Figure S1-S12.

Supplementary Table S1

Cas9 protein identity matrix

Supplementary Table S2

List of analysed Cas9 protein, gene, protein, and sgRNA sequences.

Supplementary Table S3

Position frequency matrices (PFM) of Cas9 ortholog PAM recognition

Supplementary Table S4

List of PAM interaction domain sequences and cluster information.

Supplementary Table S5

Target DNA cleavage patterns produced by Cas9 orthologs

Supplementary Table S6

List of Cas9 protein sequences used to construct the phylogenetic tree.

Supplementary Table S7

Sequences of oligonucleotides and substrates used for this study.

