## Supplemental information for "Biochemically diverse CRISPR-Cas9 orthologs"

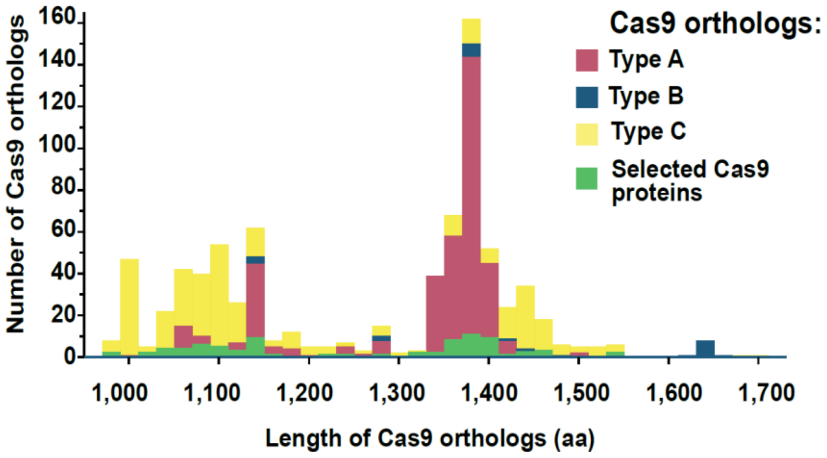


**Figure S1. Cas9 protein size distribution according subtype of CRISPR-Cas system**.

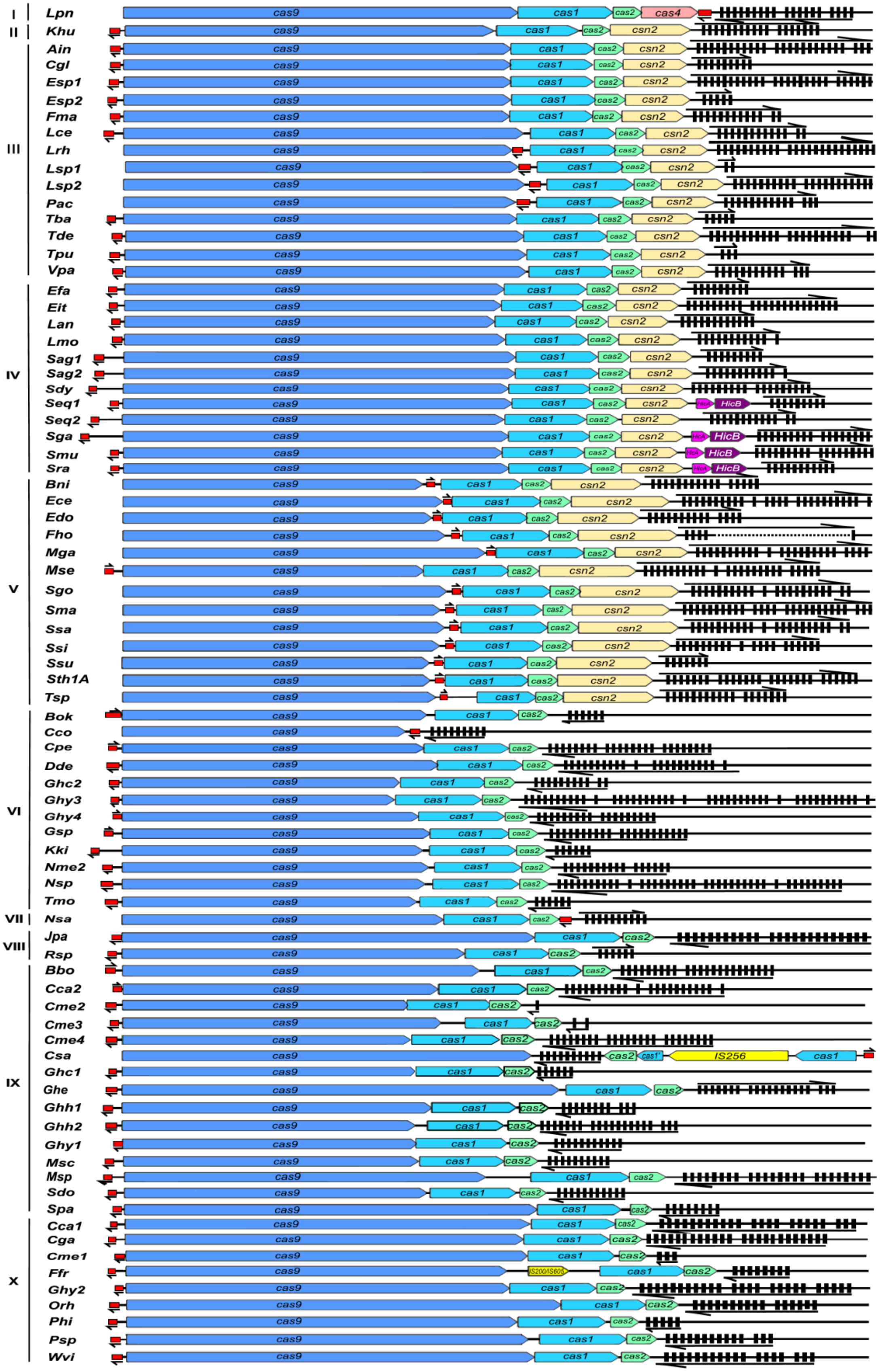

I
Lpn cas9 cas1 cas2 cas4
II
Khu cas9 cas1 cas2 csn2
III
Ain cas9 cas1 cas2 csn2
Cgl cas9 cas1 cas2 csn2
Esp1 cas9 cas1 cas2 csn2
Esp2 cas9 cas1 cas2 csn2
Fma cas9 cas1 cas2 csn2
Lce cas9 cas1 cas2 csn2
Lrh cas9 cas1 cas2 csn2
Lsp1 cas9 cas1 cas2 csn2
Lsp2 cas9 cas1 cas2 csn2
Pac cas9 cas1 cas2 csn2
Tba cas9 cas1 cas2 csn2
Tde cas9 cas1 cas2 csn2
Tpu cas9 cas1 cas2 csn2
Vpa cas9 cas1 cas2 csn2
IV
Efa cas9 cas1 cas2 csn2
Eit cas9 cas1 cas2 csn2
Lan cas9 cas1 cas2 csn2
Lmo cas9 cas1 cas2 csn2
Sag1 cas9 cas1 cas2 csn2
Sag2 cas9 cas1 cas2 csn2
Sdy cas9 cas1 cas2 csn2
Seq1 cas9 cas1 cas2 csn2 HicB
Seq2 cas9 cas1 cas2 csn2
Sga cas9 cas1 cas2 csn2 HicB
Smu cas9 cas1 cas2 csn2 HicB
Sra cas9 cas1 cas2 csn2 HicB
V
Bni cas9 cas1 cas2 csn2
Ece cas9 cas1 cas2 csn2
Edo cas9 cas1 cas2 csn2
Fho cas9 cas1 cas2 csn2
Mga cas9 cas1 cas2 csn2
Mse cas9 cas1 cas2 csn2
Sgo cas9 cas1 cas2 csn2
Sma cas9 cas1 cas2 csn2
Ssa cas9 cas1 cas2 csn2
Ssi cas9 cas1 cas2 csn2
Ssu cas9 cas1 cas2 csn2
Sth1A cas9 cas1 cas2 csn2
Tsp cas9 cas1 cas2 csn2
VI
Bok cas9 cas1 cas2
Cco cas9
Cpe cas9 cas1 cas2
Dde cas9 cas1 cas2
Ghc2 cas9 cas1 cas2
Ghy3 cas9 cas1 cas2
Ghy4 cas9 cas1 cas2
Gsp cas9 cas1 cas2
Kki cas9 cas1 cas2
Nme2 cas9 cas1 cas2
Nsp cas9 cas1 cas2
Tmo cas9 cas1 cas2
VII
Nsa cas9 cas1 cas2
VIII
Jpa cas9 cas1 cas2
Rsp cas9 cas1 cas2
IX
Bbo cas9 cas1 cas2
Cca2 cas9 cas1 cas2
Cme2 cas9 cas1 cas2
Cme3 cas9 cas1 cas2
Cme4 cas9 cas1 cas2
Csa cas9 cas2 cas1' IS256 cas1
Ghc1 cas9 cas1 cas2
Ghe cas9 cas1 cas2
Ghh1 cas9 cas1 cas2
Ghh2 cas9 cas1 cas2
Ghy1 cas9 cas1 cas2
Msc cas9 cas1 cas2
Msp cas9 cas1 cas2
Sdo cas9 cas1 cas2
Spa cas9 cas1 cas2
X
Cca1 cas9 cas1 cas2
Cga cas9 cas1 cas2
Cme1 cas9 cas1 cas2
Ffr cas9 IS200/IS605 cas1 cas2
Ghy2 cas9 cas1 cas2
Orh cas9 cas1 cas2
Phi cas9 cas1 cas2
Psp cas9 cas1 cas2
Wvi cas9 cas1 cas2

**Figure S2. Schematic representation of CRISPR loci of the Type II CRISPR-Cas systems used in this study.** Cas9 and tracrRNA encoding sequences are colored in blue and red accordingly. Predicted transcription directions of tracrRNA and CRISPR loci are indicated by arrows. Clade membership is indicated by roman numerals

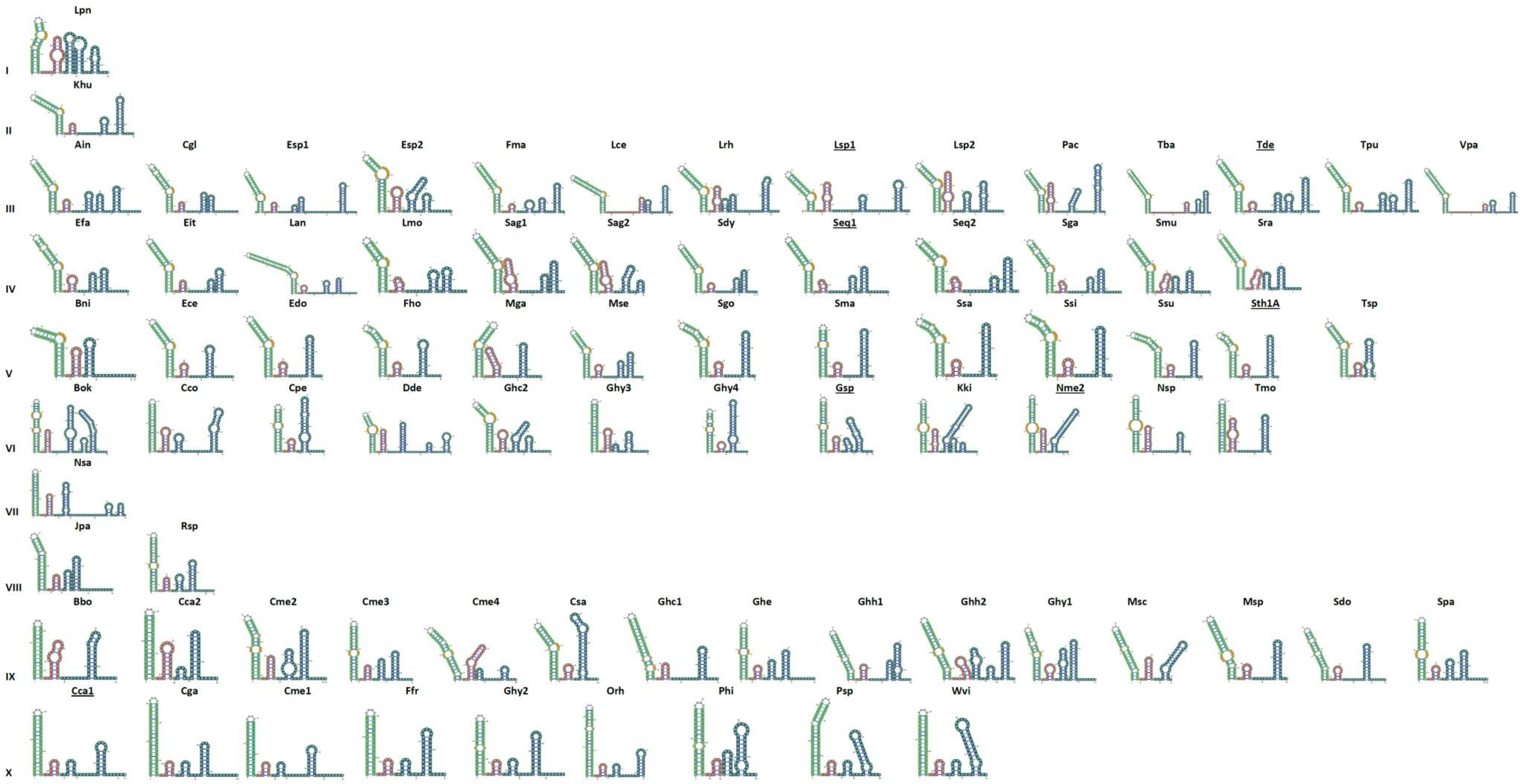


**Figure S3**. **The secondary structure diagrams of the guide RNA molecules identified for selected Cas9 orthologs**. Clade membership is indicated by roman numerals. The sgRNA structures can be classified into at least 6 groups based on structural prediction and exemplified by Tde, Seq1 (Spy-like), Sth1A, Nme2, Nsa, and Cca1 (names are underlined).

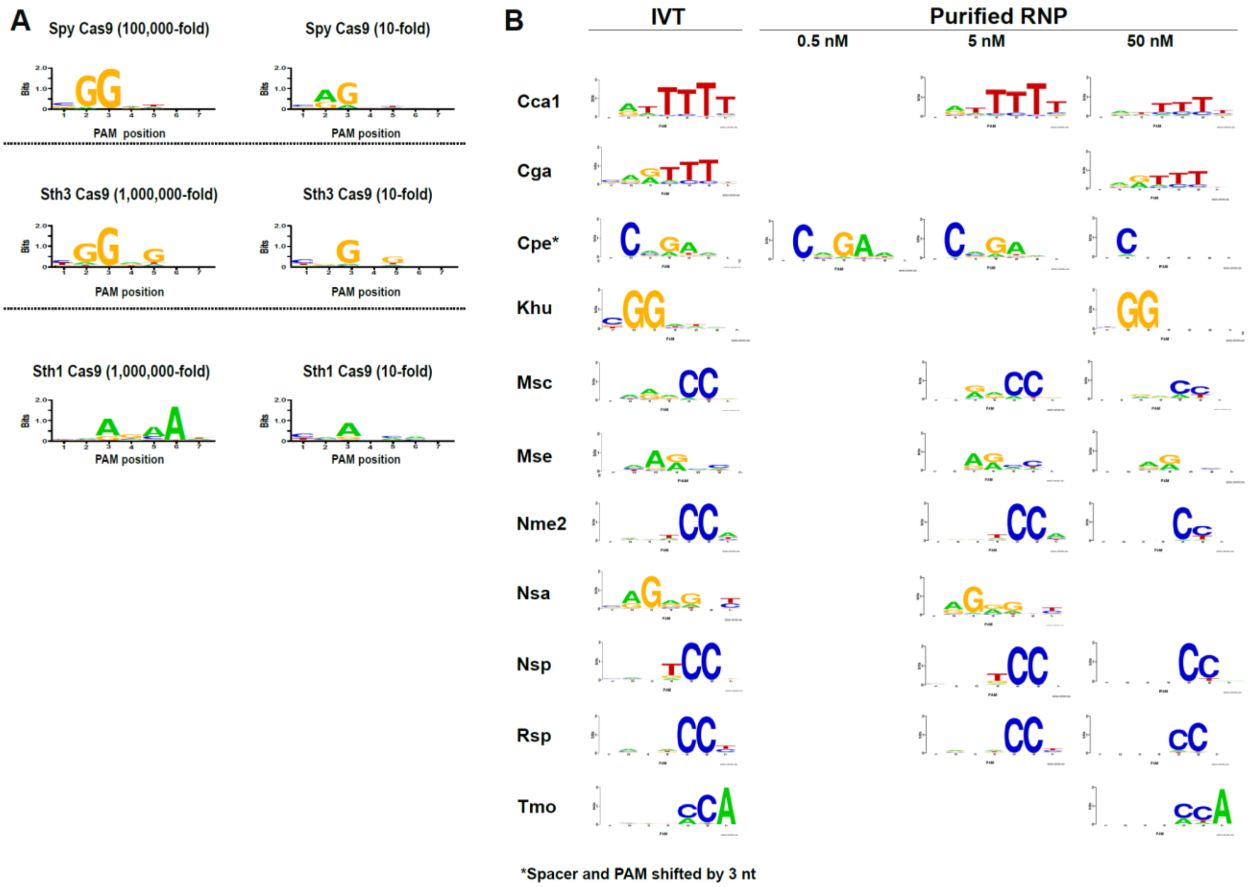


**Figure S4. Verification of PAM determination methods. (A)** PAM preferences for Spy, Sth3 and Sth1 Cas9 proteins determined using different dilutions of crude IVT RNP mixtures. **(B)** PAM confirmation using purified Cas9 proteins.

```
A) Query         C7_ID46__WP_148224960.1__ADV46720.1
Match_columns 306
No_of_seqs    35 out of 39
Neff          3.64315
Searched_HMMs 49849
Date          Fri Jan 24 13:02:17 2020
Command       hhsearch -cpu 8 -i ../results/full.a3m -d /cluster/toolkit/production/databases/hh-suite/mmcif70/pdb70 -o
../results/9327865_5.hhr -oa3m ../results/9327865_5.a3m -p 20 -Z 250 -loc -z 1 -b 1 -B 250 -ssm 2 -sc 1 -seq 1 -dbstrlen 10000 -norealign -
maxres 32000 -contxt /cluster/toolkit/production/bioprogs/tools/hh-suite-build/data/context_data.crf

No Hit                             Prob E-value P-value  Score    SS Cols Query HMM  Template HMM
  1 5CZZ_A CRISPR-associated endon  96.9 0.00033 6.7E-09   76.1  12.0  226    5-274   774-1041(1056)
  2 6KC8_A CRISPR-associated endon  96.5  0.0038 7.7E-08   68.4  15.2  200    8-274   830-1070(1083)
  3 6RJA_C AcrIIA6, CRISPR-associa  96.2  0.0088 1.8E-07   66.0  14.2  250    5-274   810-1107(1121)
  4 6JOO_A CRISPR-associated prote  87.5     1.2 2.3E-05   48.9   6.1  119    4-180   645-770 (927)
  5 5X2H_A CRISPR-associated endon  86.8     1.7 3.4E-05   47.1   6.8   79  193-274   737-822 (835)
  6 5TGY_A PS1; 4-helix bundle, co  39.6 2.1E+02  0.0043   24.1   6.6   70   54-125    31-100 (109)
...
```

```
B) Query         C8_WP_087368840.1
Match_columns 299
No_of_seqs    103 out of 122
Neff          5.0316
Searched_HMMs 49849
Date          Fri Jan 24 12:53:48 2020
Command       hhsearch -cpu 8 -i ../results/full.a3m -d /cluster/toolkit/production/databases/hh-suite/mmcif70/pdb70 -o
../results/9327865_2.hhr -oa3m ../results/9327865_2.a3m -p 20 -Z 250 -loc -z 1 -b 1 -B 250 -ssm 2 -sc 1 -seq 1 -dbstrlen 10000 -norealign -
maxres 32000 -contxt /cluster/toolkit/production/bioprogs/tools/hh-suite-build/data/context_data.crf

No Hit                             Prob E-value P-value  Score    SS Cols Query HMM  Template HMM
  1 5CZZ_A CRISPR-associated endon  99.8 8.5E-23 1.7E-27  220.1  16.6  230   46-296   768-1055(1056)
  2 6KC8_A CRISPR-associated endon  99.6   2E-17   4E-22  179.3  18.7  264    1-283   772-1071(1083)
  3 4OGE_A HNH endonuclease domain  99.1 7.2E-13 1.4E-17  144.6  10.1  226   37-293   776-1097(1101)
  4 6JOO_A CRISPR-associated prote  97.9 1.3E-06 2.7E-11   94.6  12.9  239   27-293   622-923 (927)
  5 5X2H_A CRISPR-associated endon  97.8 2.3E-06 4.7E-11   91.7  13.2  177   42-283   616-823 (835)
  6 6RJA_C AcrIIA6, CRISPR-associa  97.2 0.00038 7.6E-09   77.2  18.1  213   42-283   798-1108(1121)
  7 3BID_H UPF0339 protein NMB1088  35.9      78  0.0016   23.0   3.3   18  174-191     2-19  (64)
...
```

```
C) Query         C9_NCBI_ASWY01000120:2294253111_16
Match_columns 232
No_of_seqs    10 out of 12
Neff          3.60828
Searched_HMMs 49849
Date          Fri Jan 24 12:54:45 2020
Command       hhsearch -cpu 8 -i ../results/full.a3m -d /cluster/toolkit/production/databases/hh-suite/mmcif70/pdb70 -o
../results/9327865_3.hhr -oa3m ../results/9327865_3.a3m -p 20 -Z 250 -loc -z 1 -b 1 -B 250 -ssm 2 -sc 1 -seq 1 -dbstrlen 10000 -norealign -
maxres 32000 -contxt /cluster/toolkit/production/bioprogs/tools/hh-suite-build/data/context_data.crf

No Hit                             Prob E-value P-value  Score    SS Cols Query HMM  Template HMM
  1 6KC8_A CRISPR-associated endon  95.0   0.059 1.2E-06   56.9  10.7  174   18-211   845-1033(1083)
  2 5CZZ_A CRISPR-associated endon  94.3   0.093 1.9E-06   55.2   9.0  161   18-184   788-980 (1056)
  3 2ZDO_B Heme-degrading monooxyg  73.1      35  0.0007   23.6   7.2   61   98-160     6-68  (109)
  4 1X7V_A PA3566 protein; structu  71.9      21 0.00041   23.4   5.3   62  100-162    10-71  (99)
  5 6DS7_A Heme-degrading monooxyg  70.8      28 0.00055   23.1   5.8   64   98-162     3-66  (99)
...
```

```
D) Query         C10_WP_009541330.1__ID16
Match_columns 415
No_of_seqs    4 out of 6
Neff          1.64058
Searched_HMMs 80583
Date          Fri Jan 24 13:06:48 2020
Command       hhsearch -cpu 8 -i ../results/full.a3m -d /cluster/toolkit/production/databases/hh-suite/mmcif70/pdb70 -d
/cluster/toolkit/production/databases/hh-suite/pfama/pfama -d /cluster/toolkit/production/databases/hh-suite/NCBI_CD/NCBI_CD -o
../results/9327865_7.hhr -oa3m ../results/9327865_7.a3m -p 20 -Z 250 -loc -z 1 -b 1 -B 250 -ssm 2 -sc 1 -seq 1 -dbstrlen 10000 -norealign -
maxres 32000 -contxt /cluster/toolkit/production/bioprogs/tools/hh-suite-build/data/context_data.crf


 No Hit                             Prob E-value P-value  Score    SS Cols Query HMM  Template HMM
  1 PF13856.6 ; Gifsy-2 ; ATP-bind  71.9      12 0.00015   28.9   3.4   27  355-382    70-96  (99)
  2 PF05354.11 ; Phage_attach ; Ph  71.0      12 0.00015   30.3   3.3   27  354-380    74-100 (117)
  3 1K0H_A gpFII; twisted beta-san  64.8      15 0.00018   29.9   2.6   18  354-371    74-91  (117)
...

Cluster3 Cluster4
```

**Figure S5 HHpred results of searches using representative members from clusters 7 (A), 8 (B), 9 (C) and 10 (D).**

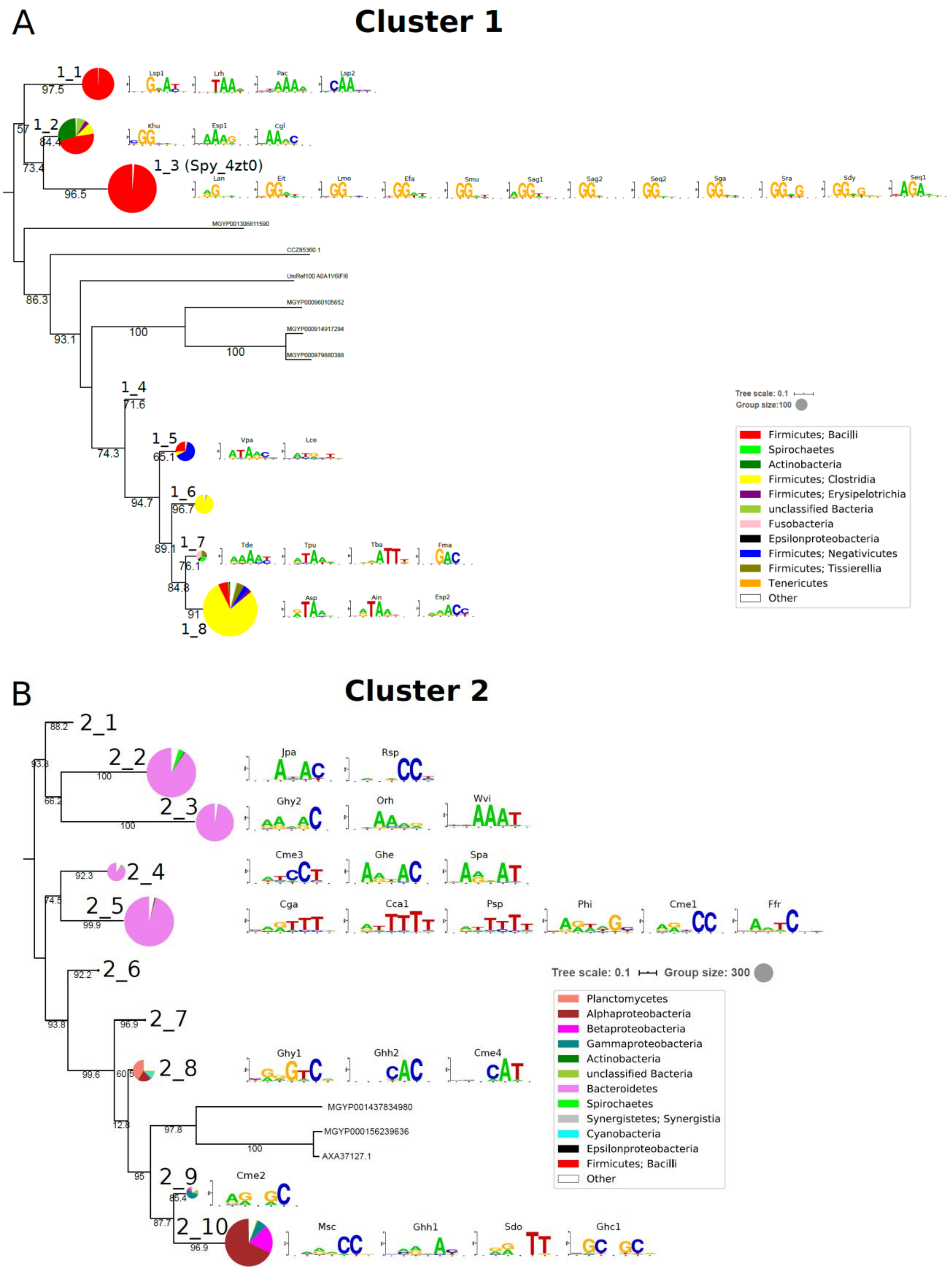
A
Cluster 1
1_1
1_2
1_3 (Spy_4zt0)
1_4
1_5
1_6
1_7
1_8
MGYP001306811590
CCZ85360.1
UniRef100 A0A1V6IFI6
MGYP000960105652
MGYP000914917294
MGYP000979880388
Tree scale: 0.1
Group size:100
Firmicutes; Bacilli
Spirochaetes
Actinobacteria
Firmicutes; Clostridia
Firmicutes; Erysipelotrichia
unclassified Bacteria
Fusobacteria
Epsilonproteobacteria
Firmicutes; Negativicutes
Firmicutes; Tissierellia
Tenericutes
Other
B
Cluster 2
2_1
2_2
2_3
2_4
2_5
2_6
2_7
2_8
2_9
2_10
Jpa
Rsp
Ghy2
Orh
Wvi
Cme3
Ghe
Spa
Cga
Cca1
Psp
Phi
Cme1
Ffr
Ghy1
Ghh2
Cme4
Cme2
Msc
Ghh1
Sdo
Ghc1
MGYP001437834980
MGYP000156239636
AXA37127.1
Tree scale: 0.1
Group size: 300
Planctomycetes
Alphaproteobacteria
Betaproteobacteria
Gammaproteobacteria
Actinobacteria
unclassified Bacteria
Bacteroidetes
Spirochaetes
Synergistetes; Synergistia
Cyanobacteria
Epsilonproteobacteria
Firmicutes; Bacilli
Other

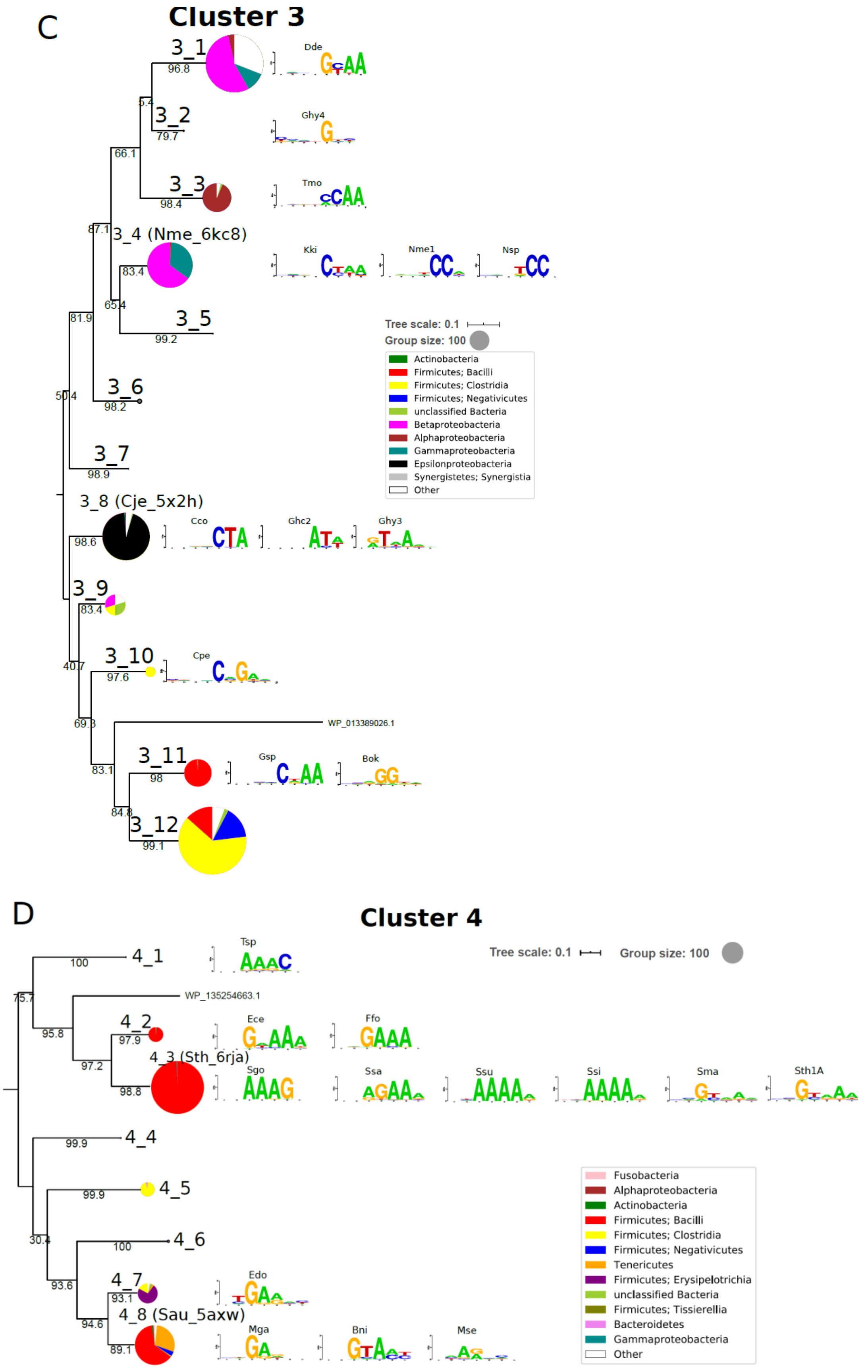
C
Cluster 3
3_1
96.8
Dde
3_2
79.7
Ghy4
5.4
66.1
3_3
98.4
Tmo
87.1
3_4 (Nme_6kc8)
83.4
Kki
Nme1
Nsp
65.4
3_5
99.2
81.9
Tree scale: 0.1
Group size: 100
Actinobacteria
Firmicutes; Bacilli
Firmicutes; Clostridia
Firmicutes; Negativicutes
unclassified Bacteria
Betaproteobacteria
Alphaproteobacteria
Gammaproteobacteria
Epsilonproteobacteria
Synergistetes; Synergistia
Other
3_6
98.2
50.4
3_7
98.9
3_8 (Cje_5x2h)
98.6
Cco
Ghc2
Ghy3
3_9
83.4
3_10
97.6
Cpe
40.7
WP_013389026.1
69.3
3_11
98
Gsp
Bok
83.1
84.8
3_12
99.1
D
Cluster 4
4_1
100
Tsp
Tree scale: 0.1
Group size: 100
75.7
WP_135254663.1
4_2
97.9
Ece
Ffo
95.8
4_3 (Sth_6rja)
97.2
98.8
Sgo
Ssa
Ssu
Ssi
Sma
Sth1A
4_4
99.9
4_5
99.9
Fusobacteria
Alphaproteobacteria
Actinobacteria
Firmicutes; Bacilli
Firmicutes; Clostridia
Firmicutes; Negativicutes
Tenericutes
Firmicutes; Erysipelotrichia
unclassified Bacteria
Firmicutes; Tissierellia
Bacteroidetes
Gammaproteobacteria
Other
4_6
100
30.4
4_7
Edo
93.6
93.1
4_8 (Sau_5axw)
94.6
89.1
Mga
Bni
Mse

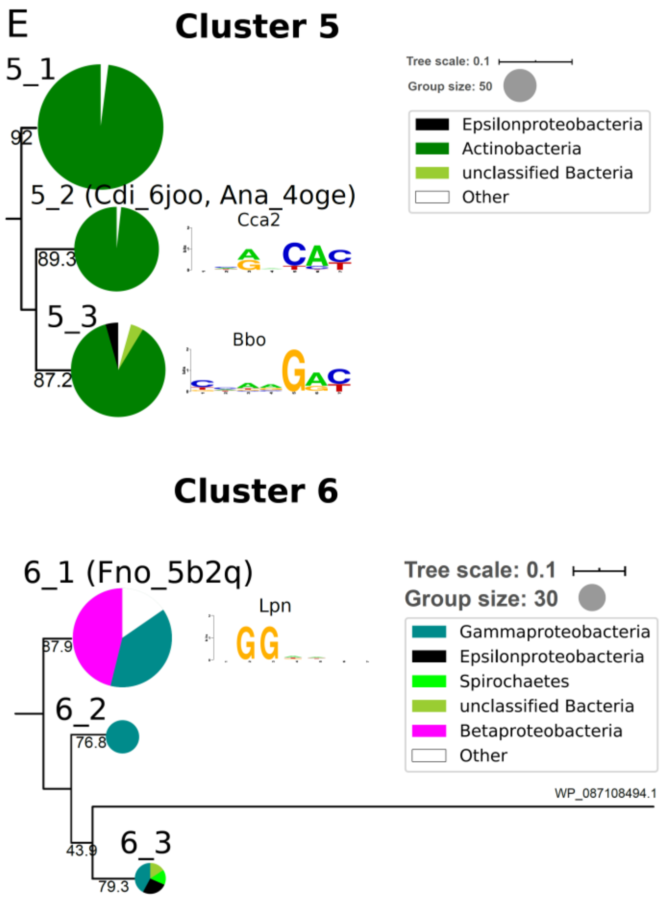


**Figure S6. Phylogenetic tree of PI domains from clusters 1 (A), 2 (B), 3 (C), 4 (D) and 5&6 (E).** SH-aLRT bootstrap values are shown at the branches.

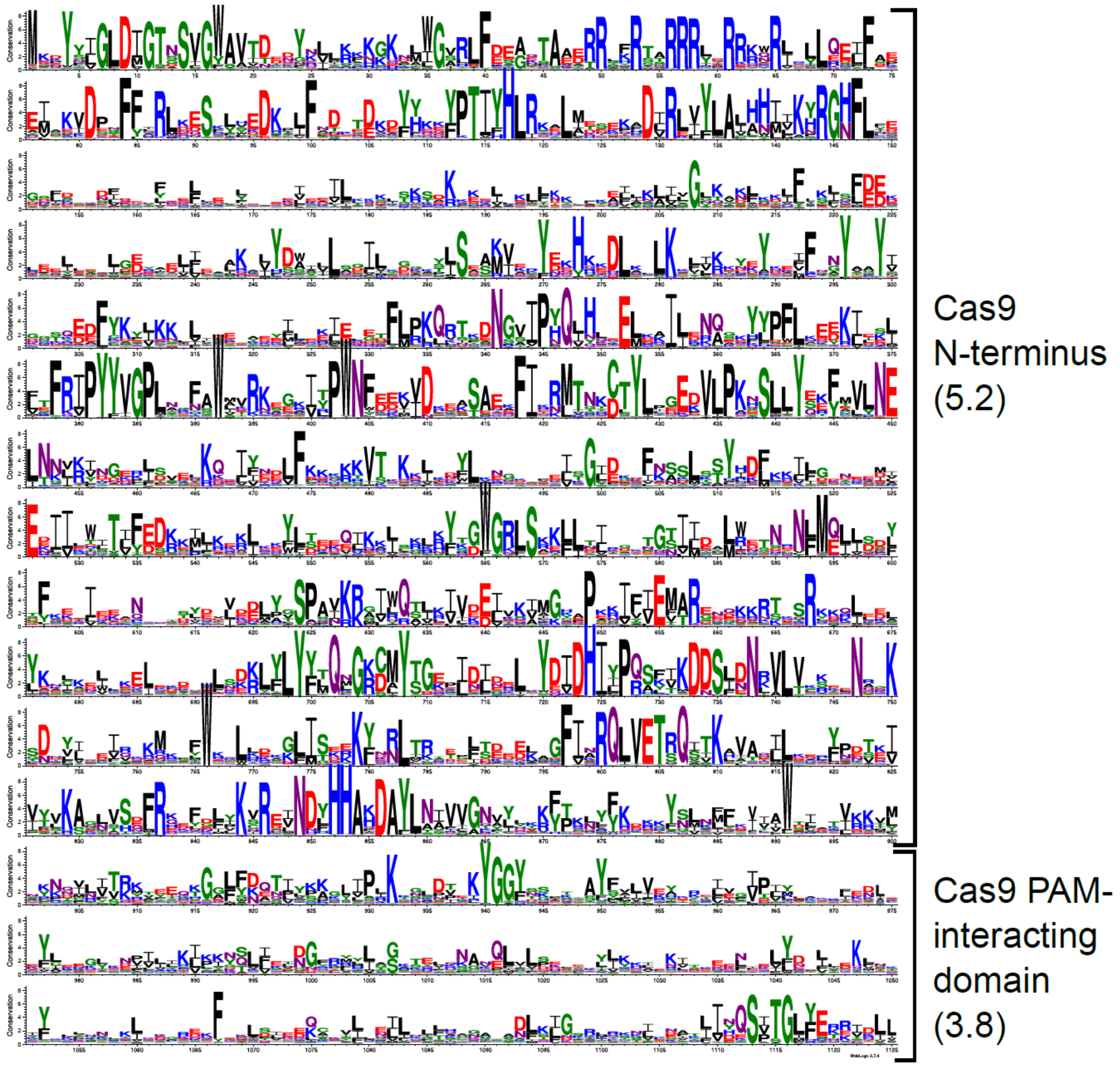


**Figure S7. Weblogo of SpyCas9 homolog sequences.** Only positions having Jalview conservation score of 1 and higher are shown. Average conservation score is shown in parenthesis. Sequences having all domains characteristic to the SpyCas9 from Cluster 1 (Supplementary table S3) were used to build MAFFT alignment.

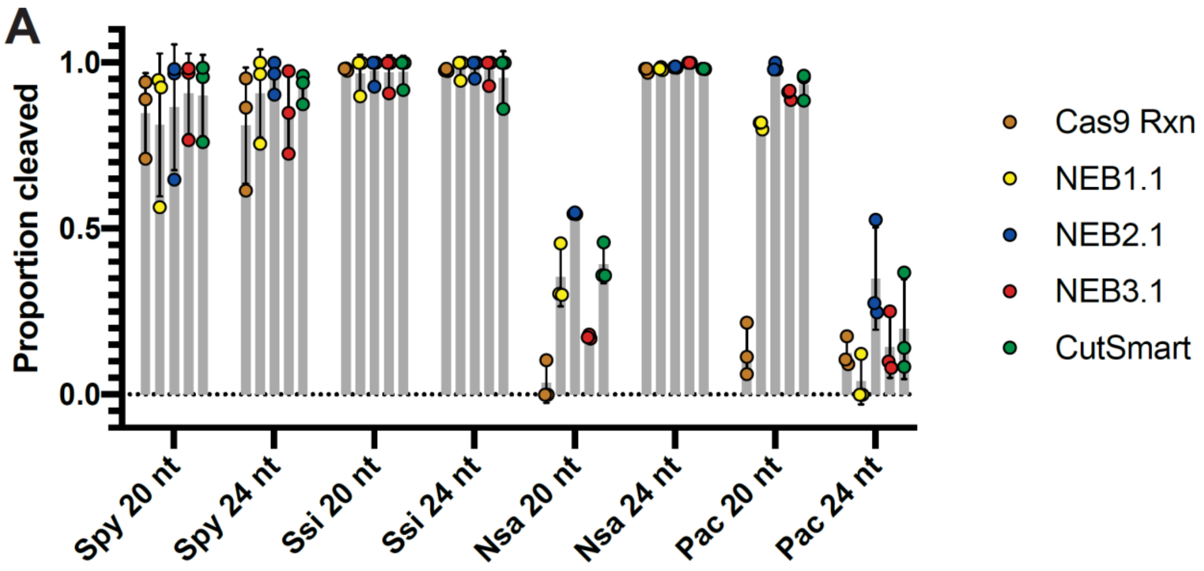

A
Proportion cleaved
1.0
0.5
0.0
Spy 20 nt
Spy 24 nt
Ssi 20 nt
Ssi 24 nt
Nsa 20 nt
Nsa 24 nt
Pac 20 nt
Pac 24 nt
Cas9 Rxn
NEB1.1
NEB2.1
NEB3.1
CutSmart

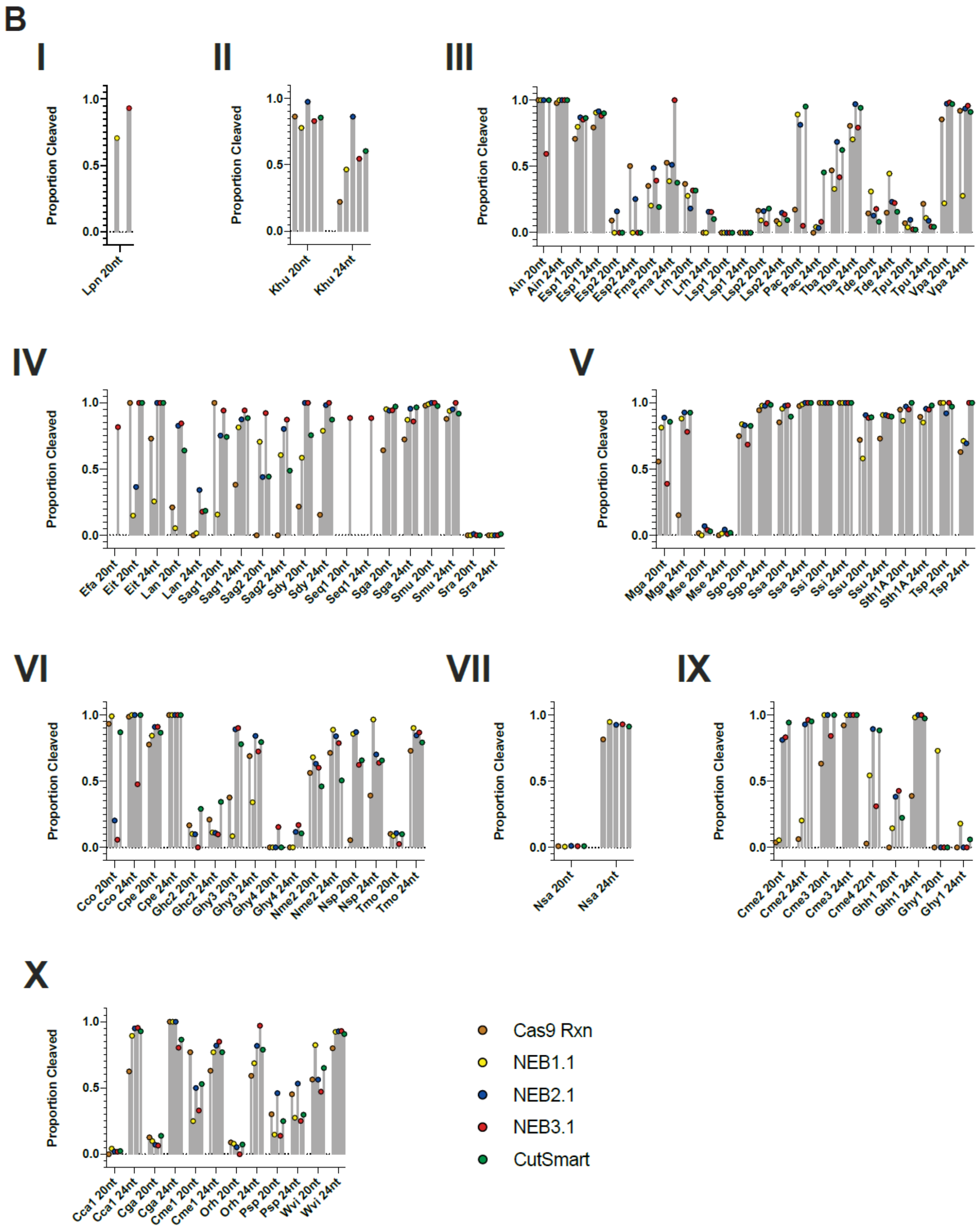

B
I
Proportion Cleaved
Lpn 20nt
II
Proportion Cleaved
Khu 20nt
Khu 24nt
III
Proportion Cleaved
Ain 20nt
Ain 24nt
Esp1 20nt
Esp1 24nt
Esp2 20nt
Esp2 24nt
Fma 20nt
Fma 24nt
Lrh 20nt
Lrh 24nt
Lsp1 20nt
Lsp1 24nt
Lsp2 20nt
Lsp2 24nt
Pac 20nt
Pac 24nt
Tba 20nt
Tba 24nt
Tde 20nt
Tde 24nt
Tpu 20nt
Tpu 24nt
Vpa 20nt
Vpa 24nt
IV
Proportion Cleaved
Efa 20nt
Eit 20nt
Eit 24nt
Lan 20nt
Lan 24nt
Sag1 20nt
Sag1 24nt
Sag2 20nt
Sag2 24nt
Sdy 20nt
Sdy 24nt
Seq1 20nt
Seq1 24nt
Sga 20nt
Sga 24nt
Smu 20nt
Smu 24nt
Sra 20nt
Sra 24nt
V
Proportion Cleaved
Mga 20nt
Mga 24nt
Mse 20nt
Mse 24nt
Sgo 20nt
Sgo 24nt
Ssa 20nt
Ssa 24nt
Ssi 20nt
Ssi 24nt
Ssu 20nt
Ssu 24nt
Sth1A 20nt
Sth1A 24nt
Tsp 20nt
Tsp 24nt
VI
Proportion Cleaved
Cco 20nt
Cco 24nt
Cpe 20nt
Cpe 24nt
Ghc2 20nt
Ghc2 24nt
Ghy3 20nt
Ghy3 24nt
Ghy4 20nt
Ghy4 24nt
Nme2 20nt
Nme2 24nt
Nsp 20nt
Nsp 24nt
Tmo 20nt
Tmo 24nt
VII
Proportion Cleaved
Nsa 20nt
Nsa 24nt
IX
Proportion Cleaved
Cme2 20nt
Cme2 24nt
Cme3 20nt
Cme3 24nt
Cme4 22nt
Ghh1 20nt
Ghh1 24nt
Ghy1 20nt
Ghy1 24nt
X
Proportion Cleaved
Cca1 20nt
Cca1 24nt
Cga 20nt
Cga 24nt
Cme1 20nt
Cme1 24nt
Orh 20nt
Orh 24nt
Psp 20nt
Psp 24nt
Wvi 20nt
Wvi 24nt
Cas9 Rxn
NEB1.1
NEB2.1
NEB3.1
CutSmart

**Figure S8. Spacer length preferences for purified Cas9 orthologs. (A)** The cleavage activity of selected Cas9 orthologs was examined using sgRNAs with 20 or 24 nt spacers. *In vitro* DNA cleavage reactions were performed in different buffers at 37ºC (see Methods). Spy and Ssi Cas9 do not have a preference for 20 or 24 nt spacers. Nsa Cas9 has a preference for 24 nt spacers whereas Pac Cas9 has a preference for 20 nt spacers. Replicate experiments are shown as points and error bars represent standard deviation. **(B)** The *in vitro* DNA cleavage activity of purified Cas9 proteins was screened using 20 and 24 nt spacers and different buffers as shown. Clade membership is indicated by roman numerals.

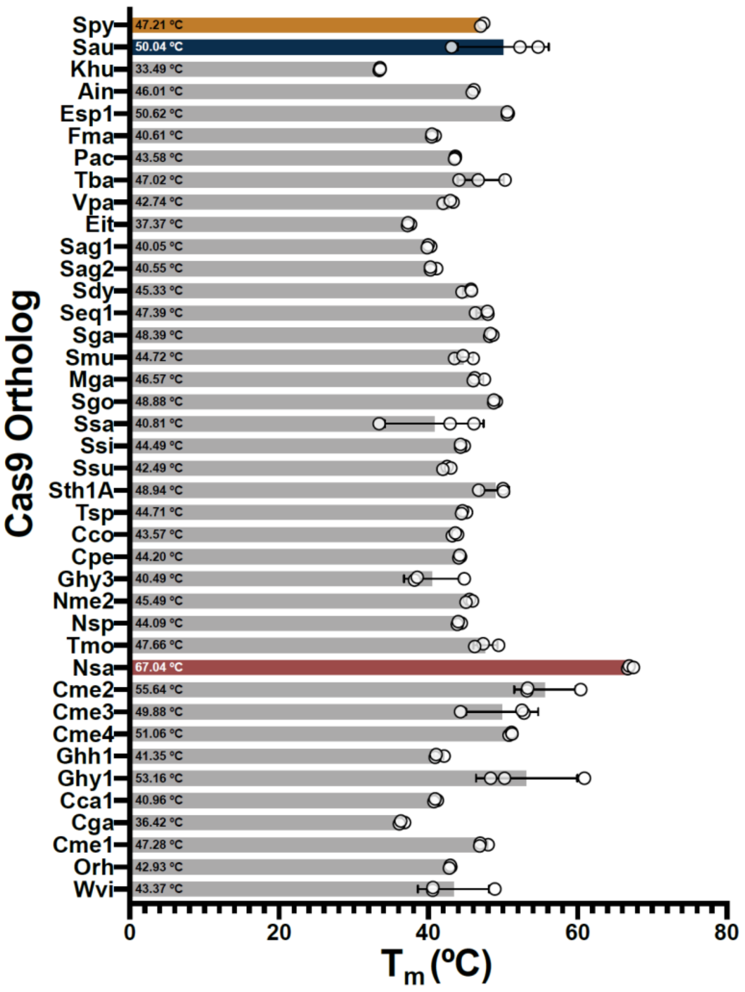


**Figure S9. Thermostability of purified Cas9 proteins**. Melting temperatures of purified Cas9 proteins was determined by nanoDSF (see Methods). Tm indicated in the histograms represent the average of 3 independent determinations plotted as open circles. Error bars indicate standard deviation.

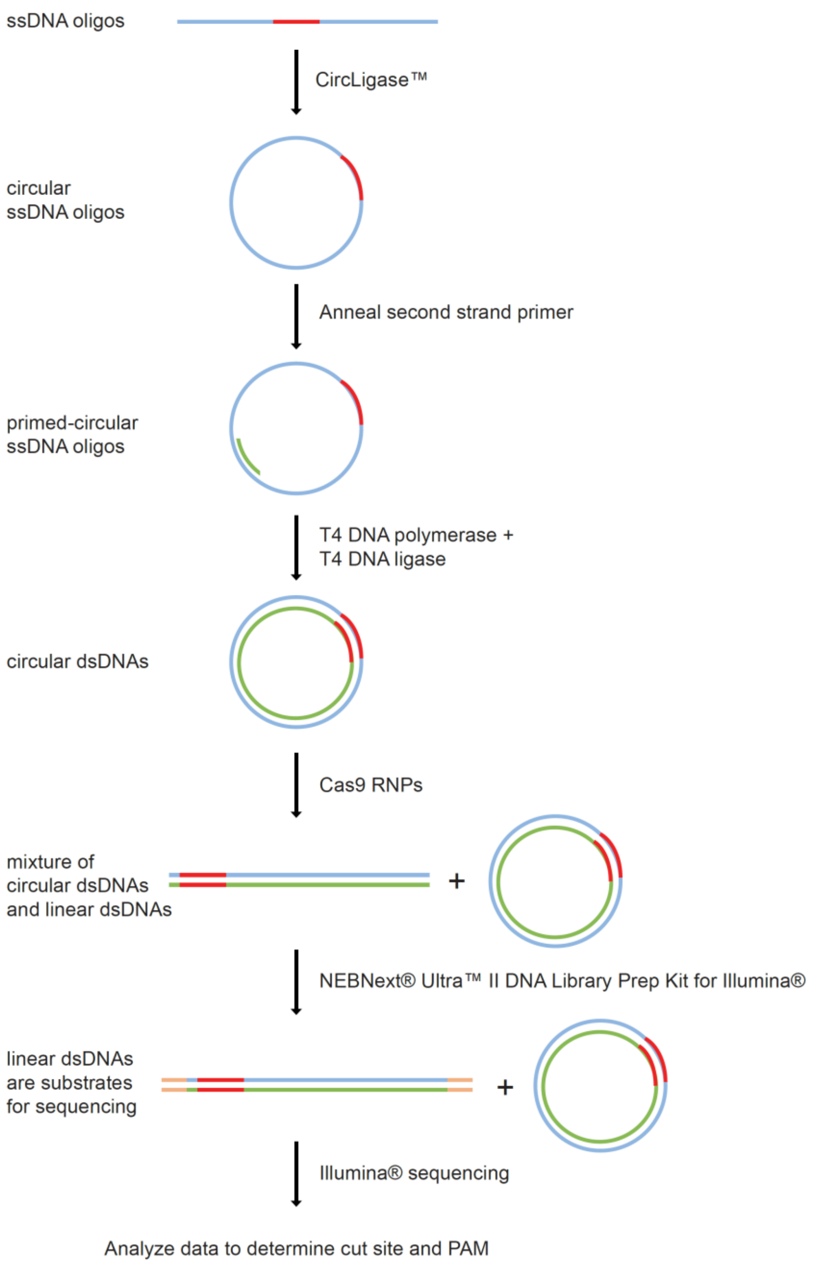


**Figure S10. PAM discovery and cleavage site mapping using purified Cas9 RNP and dsDNA minicircles.** Diagram of workflow to make small circular substrates for PAM and cut site determination assays. Pools of single stranded DNA oligonucleotides (blue) containing a 5' PO4 ends and 10 nt of randomized sequence (red) were circularized and converted to double stranded circles after annealing a second strand primer (green) and extending it with polymerase in the presence of a ligase. DNA circles are then exposed to Cas9 RNPs containing guide RNA complementary to the DNA sequence flanking the 10 nt randomized sequence region. DNA circles that are cut by the Cas9 are substrates for ligation of adapters (orange) for Illumina sequencing. Custom bioinformatic analysis was used to determine cut site pattern and PAM preference. Control reactions were carried out where each pool of circles was cut with the restriction enzyme BstXI to determine the baseline frequency of nucleotides at every position in the randomized region.

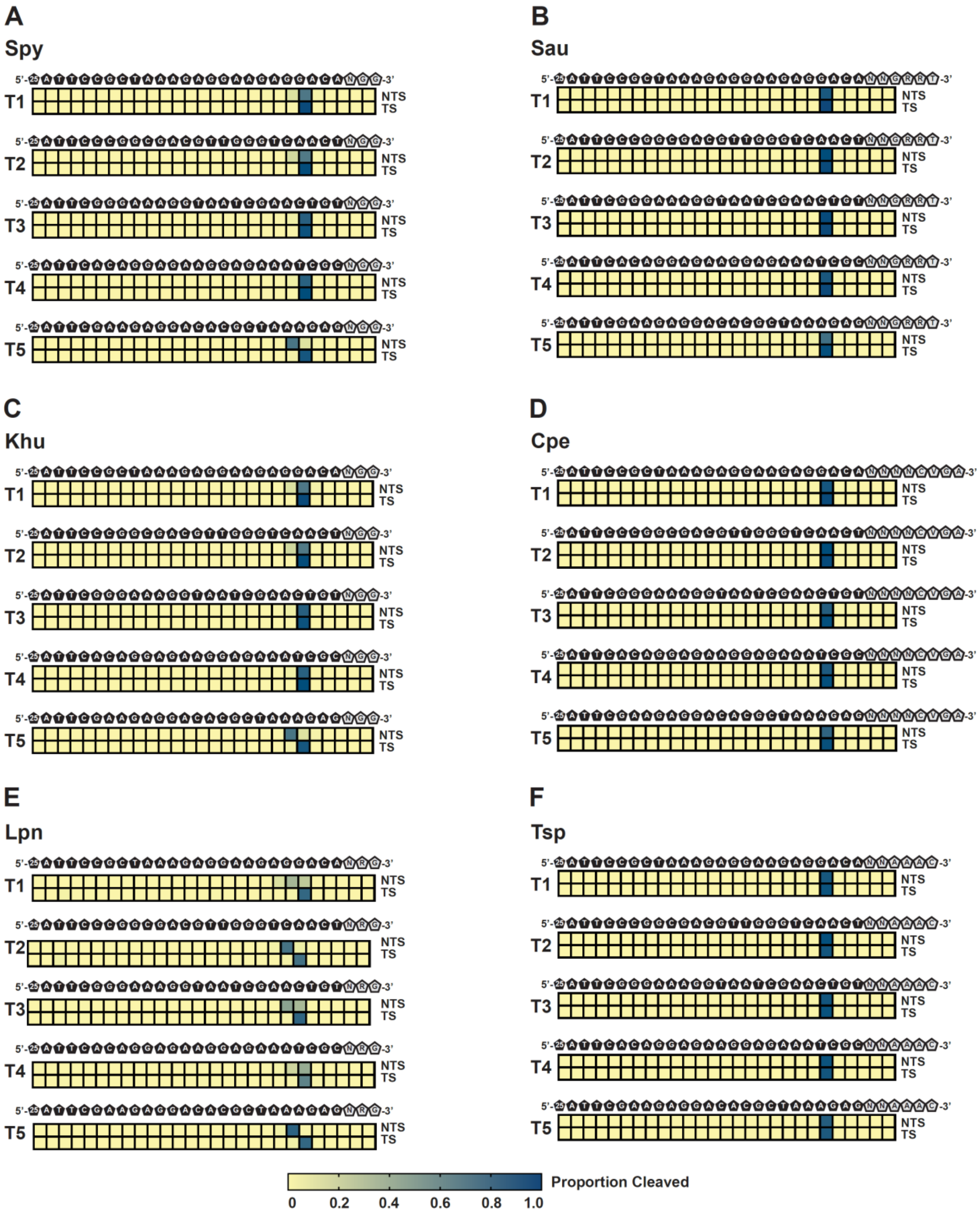

A
Spy
T1
T2
T3
T4
T5
B
Sau
C
Khu
D
Cpe
E
Lpn
F
Tsp
5'-
-3'
NTS
TS
Proportion Cleaved
0
0.2
0.4
0.6
0.8
1.0

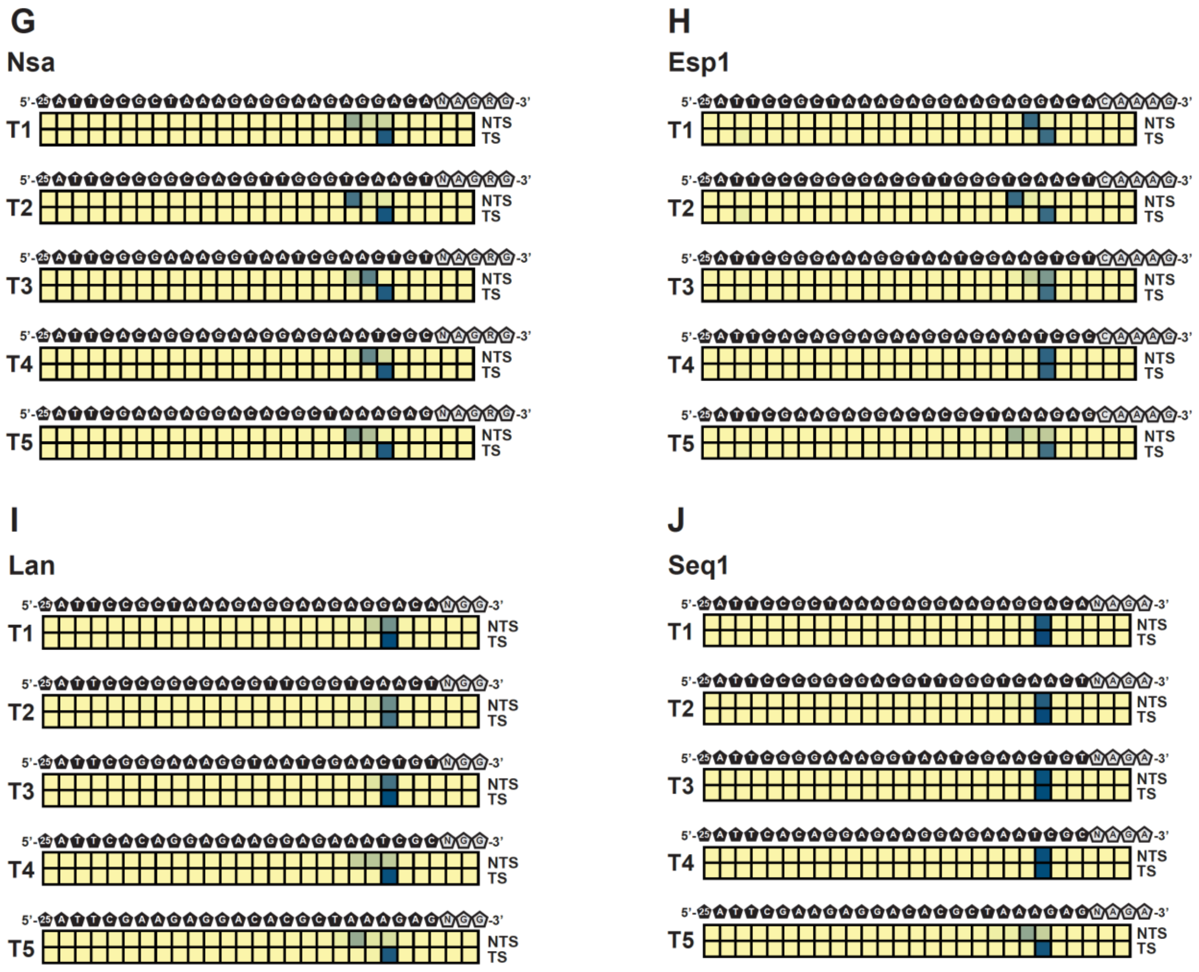


**Figure S11. Target-dependent cleavage patterns.** Diverse cleavage sites over multiple targets for a single Cas9 ortholog are depicted as heatmaps depicting mapped cleavage ends at each position in a single DNA target. Sequence of the DNA non-target strand and PAM is depicted above the heatmaps. Intensity of the blue color indicates the proportion of mapped cleavage ends. NTS indicates non-target strand; TS indicates target strand. **(A)** Spy Cas9 cleavage patterns. **(B)** Sau Cas9 cleavage patterns. **(C)** Khu Cas9 cleavage patterns. **(D)** Cpe Cas9 cleavage patterns. **(E)** Lpn cleavage patterns. **(F)** Tsp Cas9 cleavage patterns. **(G)** Nsa cleavage patterns. **(H)** Esp1 Cas9 cleavage patterns. **(I)** Lan Cas9 cleavage patterns. **(J)** Seq1 Cas9 cleavage patterns.

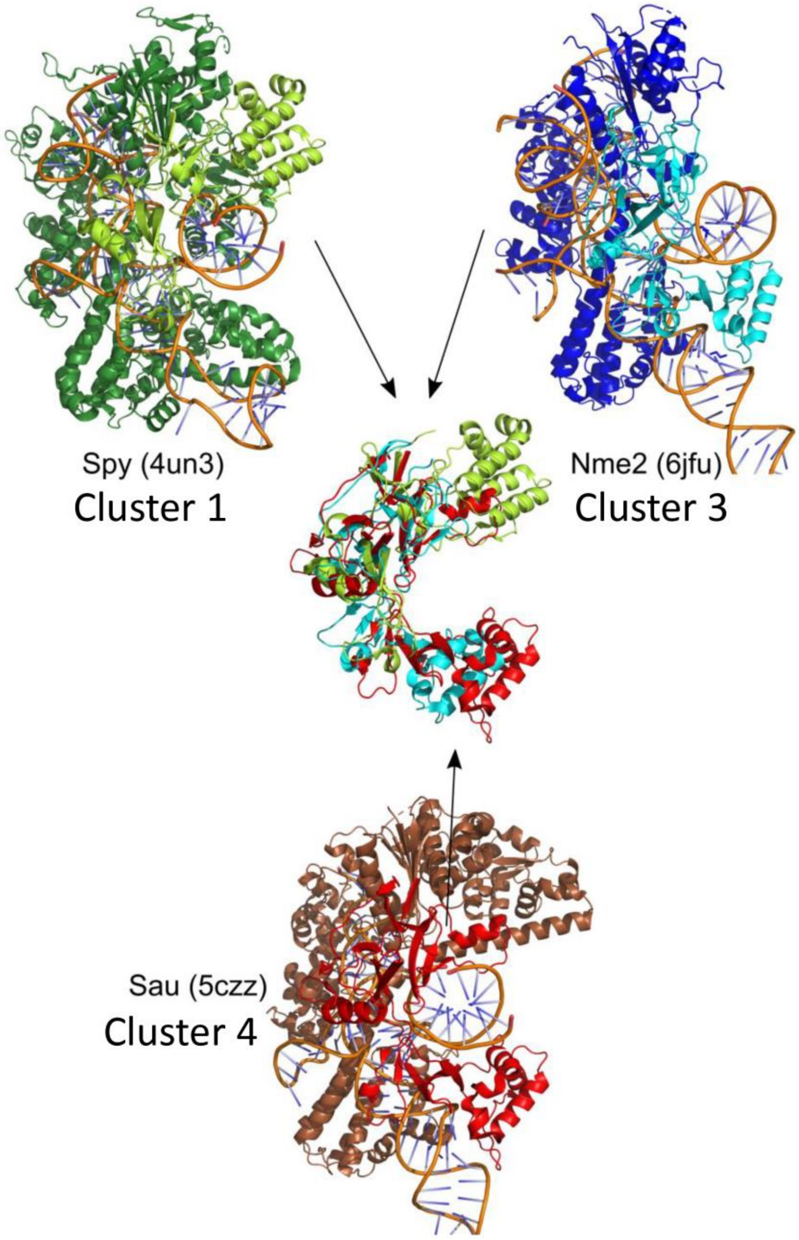


**Figure S12.** Comparison of PI domain structures from 3 major clusters (figure 2): Spy (4zto), Nmen (6kc8), Sau (5axw). There are no solved structures of Cas9 proteins which PI domain belongs to cluster cluster 2.
